# Modulating cell-free transcription electrolytically with switchable DNA triplexes

**DOI:** 10.1101/2025.04.25.650642

**Authors:** Alexander J. Speakman, Katherine E. Dunn

## Abstract

Living things self-regulate in response to various stimuli. In engineered biological systems, on-demand activation or repression of transcription underpins the design of genetic programs that control synthesis of useful products. Even electrical stimuli can control gene expression, but previous such ‘electrogenetic’ approaches have required exogenous redox mediators and complex regulatory proteins. Here we show that transcription can be modulated electrically in a cell-free system through the use of a DNA triplex as a molecular brake, which is opened or closed by means of electrolytic decomposition of water. Our DNA constructs comprise a double-stranded region with a single-stranded tail that folds over to form a triplex, where this construct is made using a customised synthesis protocol. We tested 48 triplex-forming constructs and their corresponding controls, and in numerous cases the triplex successfully inhibited gene expression in a controllable manner. Our approach, Electrically Directed Gene Expression (‘EDGE’), could be used to modulate in vitro synthesis of RNA therapeutics and also has applications in smart biomaterials, where the RNA transcript would act as a signal for a change of state. To test EDGE, we produced customised 96-well plates and control electronics, demonstrating production of experimental hardware via 3D-printing and in-house manufacturing.

## [Main text]

The ability to regulate gene expression is of vital importance in biological systems, most evidently *in vivo* for processes such as homeostasis [1], but also *in vitro* when using cell-free systems to manufacture valuable biological products, such as therapeutic proteins or mRNA vaccines [2] [3]. External regulation of gene expression in cell-free systems can enable greater control over the quantity of expressed product, the timing of expression, or the conditions under which expression occurs [4] [5]. There are many methods of controlling gene expression i*n vitro*, including optogenetic systems that use light as a stimulus to turn expression on or off [6] [7] and chemical control systems such as the Lac operon [8] or TET on/off systems [9] that utilise the addition of an exogenous chemical (lactose or tetracycline respectively) to stimulate or halt gene expression. These strategies can be considered expandable, via the use of a variety of regulatory proteins that respond to separate triggers, including light of different wavelengths or different chemical stimuli.

As the market for biologics expands there will be an increasing need for diverse, programmable control mechanisms for cell-free synthesis of high-value products, driving a search for alternatives to conventional chemical and optogenetic methods. It will be valuable to consider how electrical stimuli could be used to drive changes in gene expression, an area known as electrogenetics. In one previous electrogenetic approach, electrochemical oxidation of ethanol into aldehyde was used to control CHO cells engineered to express aldehyde induced transactivator proteins [10], while in another platform reversible reduction and oxidation of pyocyanin controlled the endogenous *E. coli* SoxR pathway to turn gene expression on and off as needed [11]. Such electrogenetic systems are limited by the need for exogenous redox mediators and complex regulatory proteins, with some approaches also suffering from the disadvantage of requiring cells to be grown in the presence of potentially toxic chemicals such as ethanol and aldehyde.

With conventional non-electrical methods for control of gene expression, it has already been seen that using nucleic acid switches reduces the dependence on regulatory proteins. For example, modulation of RNA cleaving can be used to control expression in mammalian cells [12] [13] and rationally designed strand displacement systems have been deployed to direct translation [14]. It is known that non-canonical DNA structures have a role in the natural regulation and dysfunction of gene expression [15] [16] [17] [18], while triplex-forming oligonucleotides can be used to target genes via binding to specific sequences within a duplex [19]. With a cell-free system, it has even been shown that a strand displacement reaction can be used to release a triplex that would otherwise inhibit the action of an RNA polymerase [20]. By exploiting pH-sensitive DNA triplexes [21], we have now developed a method that we call Electrically Directed Gene Expression (EDGE), which employs DNA triplexes as electrically controllable switches for activating gene expression in a cell-free medium, providing a direct bridge between an electrical system and a genetic program. Our primary goal was to develop a transcriptional regulation strategy that offered an advantage over alternative approaches because it eliminated the need to use regulatory proteins or potentially toxic additives such as inducers or mediators. We also aimed for our system to be as simple as possible, with the minimum number of active elements.

### Mechanism for DNA triplex-mediated electrolytic control of transcription

The DNA construct we developed for EDGE comprises a linear double-stranded (ds) segment and a long overhanging single-stranded (ss) tail - the triplex-forming region (TFR) (Fig. 1a). The double-stranded section contains a promoter, a downstream triplex-binding region (TBR) to which the tail binds, and finally a product coding region. We chose the fluorogenic RNA aptamer iSpinach [22] as a product, to enable transcription output to be quantified via fluorescence measurements without the need for translation [23], and we selected the T7 cell-free expression system due to its accessibility and compatibility with iSpinach. In early experiments, we attempted to use the Expressway system but found this was not compatible with iSpinach (see supplementary note) and opted for a customised formulation (Methods). In our experiments, when the triplex is closed, the action of the T7 polymerase is inhibited and transcription is slowed or halted. The triplex results from formation of additional hydrogen bonds called Hoogsteen interactions between ssDNA and dsDNA regions with appropriate sequences [24] (Fig. 1b). Depending on the sequence and directionality of the ssDNA strand with regard to the dsDNA, different base triplet combinations are possible [25]. EDGE utilises polypyrimidine, parallel, major groove triplexes that can form T•AT and C^+^•GC triplets (ssDNA base • dsDNA pair). Here, C^+^ refers to protonated cytosine, which is produced under acidic conditions [26], providing the required pH-dependence for our application. The pH-sensitivity is tuneable based on the base composition of the triplex, because including more cytosine bases requires greater protonation and lowers the pH level at which the triplex is stable. Idili *et al.* [21] showed that triplex-based switches composed entirely of T•AT triplets would only undergo an unfolding transition at around pH10 while a switch where some of the triplets were C^+^•GC would unfold at more basic pHs; a 60% T•AT switch is fully unfolded at pH9.5, while a 20% T•AT switch unfolds below pH7.

**Figure 1:**
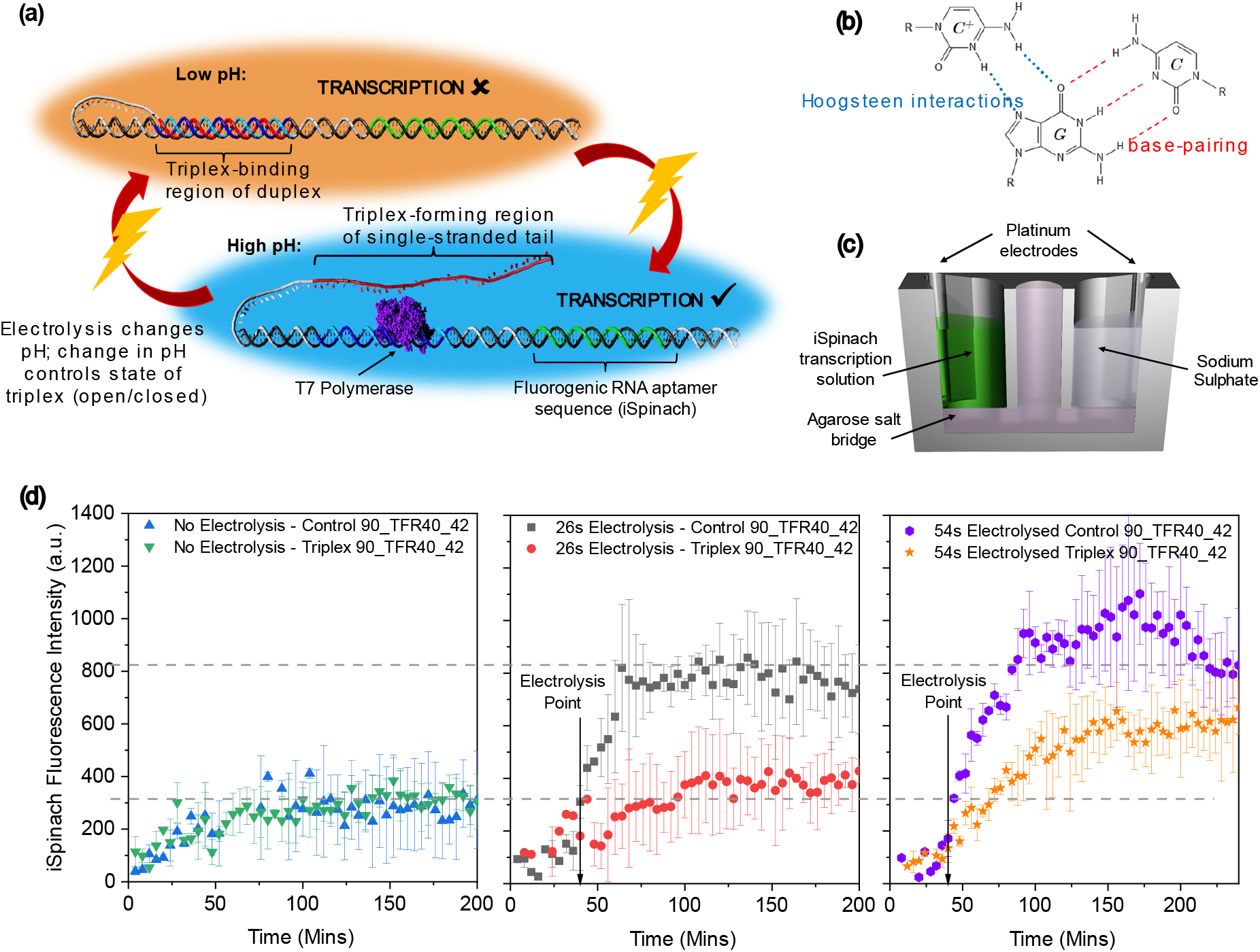
Overview of our Electrically Directed Gene Expression (EDGE) technology. (a) Our DNA construct consists of a double-stranded domain (encoding the promoter, gene of interest etc) and a single-stranded tail that can fold over to bind with a segment of the double-stranded region, forming a triplex. The sequences are designed such that triplex formation is pH-dependent. Electrolytic decomposition of water is used to change pH and switch triplex state. (b) Illustration of the interactions underpinning the formation of a single triplet as part of a three-stranded complex, where the presence of the protonated cytosine drives the pH-dependence. (c) Schematic cross-sectional view of the electrical setup. The sacrificial well contains sodium sulphate while the other contains the cell-free transcription medium. The wells are connected electrically by an agarose-based salt bridge. A constant current is applied via platinum electrodes. (d) iSpinach fluorescence during transcription, observed from a mixture containing a representative triplex construct and another with the corresponding control construct (not triplex-forming). Buffer = 1X 1% AceOTB (see Methods). Data is presented for three scenarios: (1) no electrolysis is performed (left panel), (2) mixtures are exposed to 26 seconds of current (middle panel) (3) electrolysis duration is increased to 54 seconds (right panel). Grey dotted lines are guides to the eye. Error bars = standard deviation (*N*=3).

Previously published work has demonstrated the reversible electrical control of pH-dependent triplex formation through the use of a quinhydrone redox system [27]. In contrast, we use the electrolytic decomposition of water to achieve a pH shift [28]. Reduction at the cathode produces hydrogen and OH^-^ ions, while oxidation at the anode produces oxygen and H^+^ ions [29], and we can use this to shift the pH of our cell-free solution up or down depending on the polarity of current applied. Consequently, we could control the state of our DNA triplex using only electrodes and a current source, with appropriately designed hardware and wetware. A typical setup featured two wells, one of which contained the iSpinach transcription solution with our chimeric ss/ds/triplex DNA construct while the other contained sodium sulphate as a conductive electrolyte (Fig. 1c). The two wells were isolated to prevent diffusion but we completed the circuit via an agarose-based salt bridge.

In a representative experiment, we observed minimal iSpinach fluorescence from both our triplex-forming construct and a non-triplex-forming control (Fig. 1d). Here, the triplex-forming region was 40 base pairs in length and the triplex composition was 90% T•AT, while the inactive linker section of the tail was 42 nucleotides long. In this and all our other experiments, the double-stranded segments of the control construct were identical to that of the corresponding triplex-forming constructs (defined by the TFR length and TBR composition), while the single-stranded tail of the control construct was designed such that it was incapable of forming a triplex. Throughout our study, it was vital to compare the performance of triplex-forming constructs with that of non-triplex-forming controls, as transcription is inherently pH-dependent and we wish to demonstrate an ability to control or modulate this natural behaviour using a synthetic molecular switch.

In the absence of electrolysis, very little iSpinach was produced from either construct because the pH was too low for optimal transcription (Fig. 1d, left panel). In a parallel experiment, we measured iSpinach fluorescence after 26 seconds of constant current electrolysis. Here the pH was estimated to be 7.25 (calibration described later in the manuscript), closer to the optimum required for transcription, and we saw that iSpinach levels rose dramatically after electrolysis of the solution containing the control construct (Fig. 1d, central panel). In comparison, minimal activation was apparent from our triplex-forming construct, indicating that the triplex was still closed and inhibiting transcription. In a third experiment, we applied an electrolysing current for 54 seconds, bringing the pH to approximately 7.5. Here, the amount of iSpinach produced by the control construct was slightly greater than that seen in the 26 second case, but the triplex-forming construct displayed considerably greater transcription activity than for the previous experiment. Consequently, we conclude that after 54 seconds of electrolysis the pH has changed sufficiently to disrupt the inhibitive effect of the triplex and enable transcription to occur.

Developing our electrogenetic platform required us to validate triplex behaviour, identify a buffer compatible with both triplex formation and transcription, and develop a pathway for producing an unusual DNA construct with both ss and ds regions. To shed light on the significance of different features of EDGE constructs we made 48 design variants (with their associated controls), which also required us to produce customised hardware for high-throughput measurements, with precisely calibrated current pulses for electrolysis.

### Validating triplexes and transcription buffers

We began by using fluorimetry to confirm that electrolytic pH shifting could be used to form or destabilise triplexes. In this experiment, we used a pair of two cuvettes, each with an inserted electrode and connected by a salt bridge, placed in a fluorimeter with a customised 3D-printed holder (Fig. 2a). We added a stirrer motor to one of the cuvettes, in which we placed a solution of a simplified triplex construct that was labelled with a Cy5 fluorophore and a quencher such that fluorescence was suppressed when the 10-nucleotide-long triplex was closed. Starting from a low pH and with the electrode in the triplex cuvette as the cathode, electrolysis induced an increase in pH that caused fluorescence to increase sharply, indicating that the triplex structure had been destabilised (Fig. 2b). Reversing the polarity and electrolysing again reversed the pH change, causing a similarly sharp drop in fluorescence, indicating that the triplex had been reformed. This confirmed that electrolytic decomposition of water could be used to control triplex formation in a reversible manner. We were able to cycle the triplex between open and closed states, although the signal decreased with cycle number, an effect that we attribute to degradation of the fluorophore.

**Figure 2:**
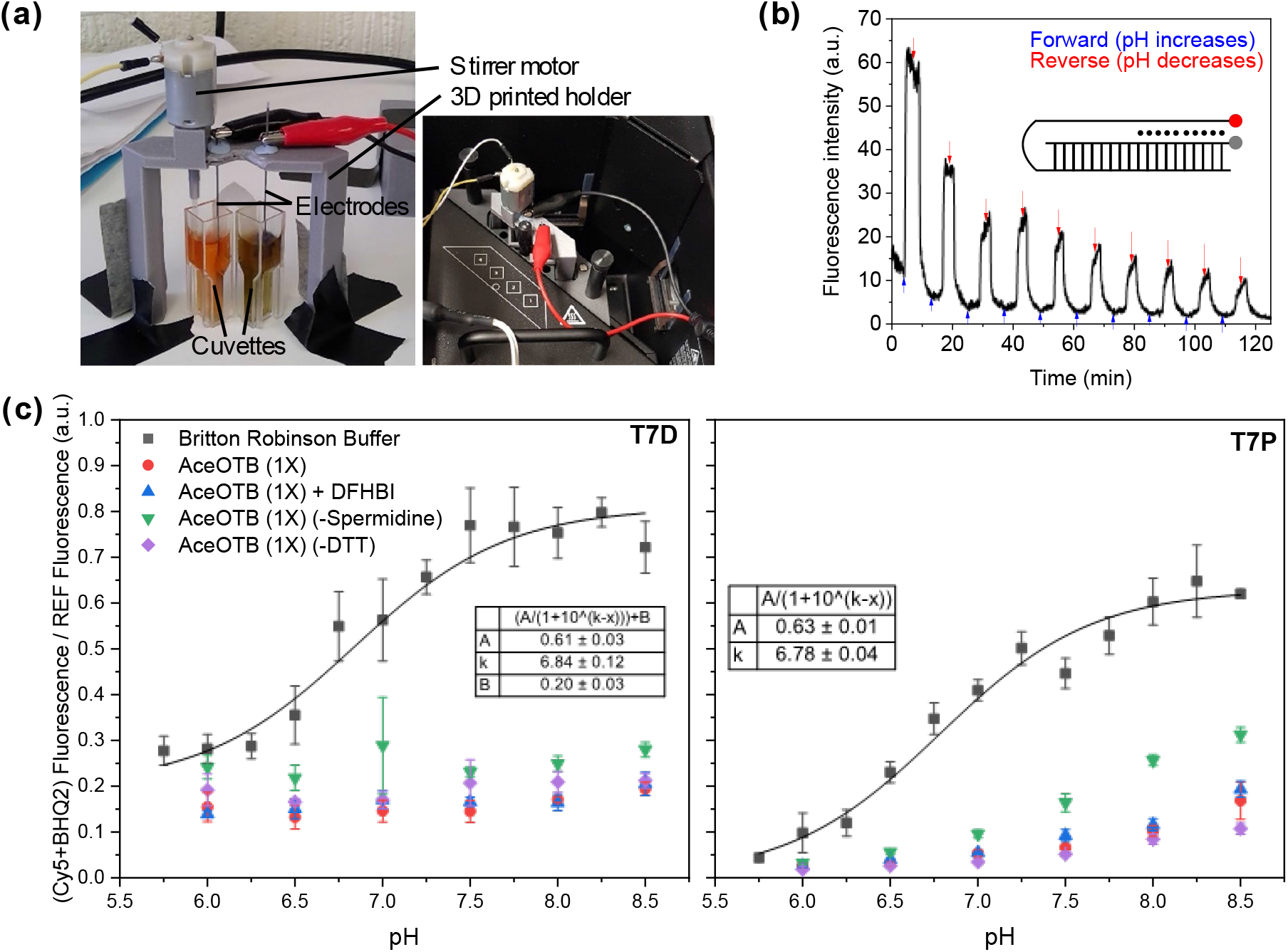
Electrolysis, pH and DNA triplexes. (a) Left: 3D-printed mount holding stirrer and graphite electrodes in a pair of cuvettes, each containing 2mL of 1M sodium sulphate with 20μL universal indicator, with a 5V source used for electrolysis. Before electrolysis, both samples were yellow. Image shows post electrolysis states: Murky green (cuvette on right) = higher pH, orange/red (cuvette on left) = lower pH. Right: the cuvette/motor/electrode assembly installed in a fluorimeter. (b) A short ten-nucleotide triplex (TFR10 construct) is cycled between open and closed states by electrolytic control of pH, using the setup in the right-hand photo of part (a). The triplex is labelled with a Cy5 fluorophore and a quencher (inset – red circle = Cy5, black circle = fluorophore). Fluorescence is low when the triplex is closed and increases upon opening. Electrolysis with ‘forward’ polarity increases pH, ‘reverse’ polarity decreases pH. (c) Formation dynamics of more complex triplexes in various buffers as a function of pH, in the absence of electrolysis, measured using Cy5-quencher-labelled triplexes. T7D refers to T7D_TFR20_54 and T7P refers to T7P_TFR40_N9. Both triplexes display the expected transition in Britton Robinson buffer, with a midpoint around pH8.6 but are more stable in AceOTB even in the absence of spermidine or DTT. Data is presented as a fraction of the signal for a non-triplex forming construct. Here, AceOTB=1X 100% variant (see Methods).

In the EDGE constructs, a long single-stranded linker was required for the TFR to reach the TBR. However, it is known that the length of the linker affects the stability and pKa of DNA triplexes [30] due to the entropic penalty of a longer loop. Thus we needed to confirm that our EDGE constructs could still form triplexes in spite of the length of the linker domain. We examined two designs – T7P, in which the triplex formed within the promoter domain (tested with a 40nt triplex and 9nt linker), and T7D, in which the triplex formed downstream of the promoter (tested with a 20nt triplex and 54nt linker). As before, the constituent oligonucleotides were labelled with a Cy5 fluorophore and a black hole quencher such that the fluorescence would drop when the fluorophore and quencher were brought into proximity with each other upon triplex formation. In this case we also performed control measurements on a duplex labelled with Cy5 and no quencher. Our first experiments (Fig. 2c,d) were performed using Britton Robinson buffer adjusted to different pH levels [31]. The results demonstrated that both T7D and T7P constructs were capable of forming triplexes, giving characteristic sigmoidal curves with similar transition midpoints (around pH6.8). The width of the transition was similar although the T7D curve was offset vertically. For the purposes of the fluorescence experiments the constructs were truncated and the coding region was omitted.

We proceeded to test triplex formation in the buffer to be used for transcription (AceOTB – Acetate Optimised Transcription Buffer). Compared to Britton Robinson buffer, in which we observe a definitive destabilisation of triplexes and increase in fluorescence as pH increases, the same constructs in AceOTB do not appear to open in the same manner and are remarkably stable even at pH8.5, even with the removal of specific components such as spermidine and DTT. At first this appears to suggest that we would normally see no expression from EDGE constructs because the triplex would be closed under any practically achievable pH condition, preventing the polymerase from accessing the promoter. However, RNA polymerase can exert substantial forces [32] and could potentially force open a weakened triplex. This should give rise to a balance between the pH-dependent strength of the triplex and the ability of the polymerase to force it open. We hypothesise that this phenomenon enables transcription to occur even under conditions that would normally keep the triplex closed and we discuss this theory in greater detail later in the manuscript. The possibility of protein machines unwrapping DNA structures is not unknown; it has been observed that ribosomes can perform translation from an mRNA strand that is initially embedded in part in a DNA origami structure [33].

### Production and testing of EDGE constructs

For EDGE, we required unusual chimeric DNA constructs with both double and single-stranded sections, necessitating a novel synthesis method (Fig. 3a). We produced our EDGE constructs via hybridisation of two complementary strands, where one had a long overhang (up to 94nt) that acted as the triplex-forming tail. As the complexity and length of the sequences used were too difficult for commercial ssDNA oligonucleotide synthesis, we embedded the sequences in plasmids, which we amplified by PCR to form linear dsDNA constructs, before using asymmetric PCR (aPCR) [34] [35] to generate each strand of the constructs separately. As the dsDNA templates would be capable of transcription if left in solution, the initial PCR reaction was performed using with one biotinylated primer enabling removal of excess dsDNA through a streptavidin pulldown [36]. We optimised our method using representative construct designs and further purified the single-stranded components by extracting them from an agarose gel and hybridising them as necessary. We then performed further electrophoresis to confirm that the aPCR and hybridisation steps had occurred as expected (Fig. 3b and Extended Data Figure 2). In all cases, for the lanes containing one strand we obtained a well-defined band of the expected size. In some cases, additional weak bands were visible (e.g. T7P-bottom strand), which we associate with the formation of multi-strand complexes and/or secondary structure motifs. In the lanes containing hybridisation products, there is usually one band at the same level as one of the two constituent strands due to imperfect stoichiometry. For triplex-forming constructs, the hybridisation product may also give rise to two bands – one in which the triplex is closed and one in which it is open. It is also possible for faint bands to arise due to the formation of a small number of multimeric complexes.

**Figure 3:**
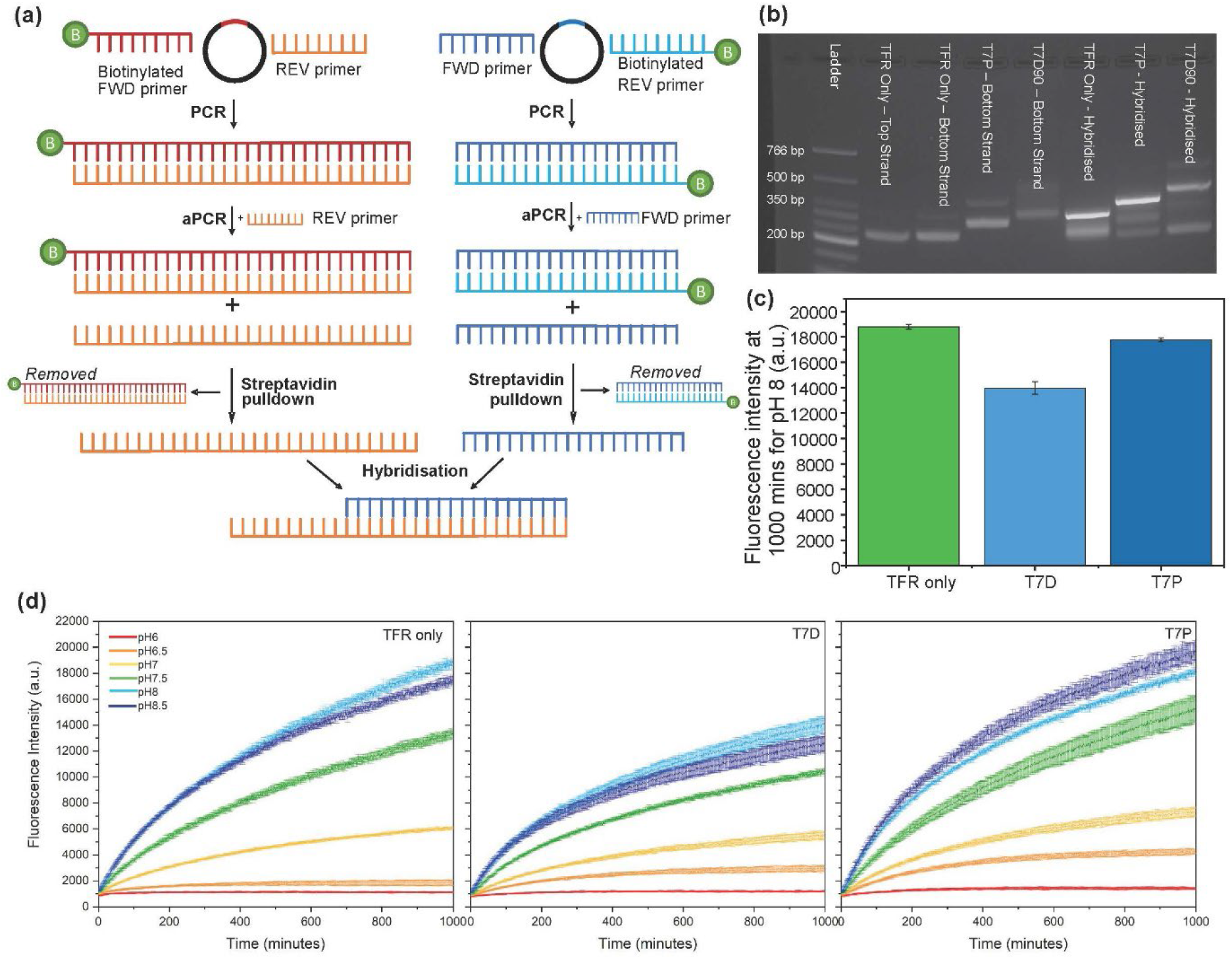
triplex-based constructs – synthesis and use for cell-free transcription. (a) aPCR workflow. The sequences for the EDGE construct with and without tail were synthesised as dsDNA plasmids (shown as circles). PCR step: We then amplified the relevant sections of the plasmids to form a linear dsDNA template. Here, one of the primers was biotinylated, depending on which strand (‘top’ – blue, right-hand side; ‘bottom’ – orange, left-hand side) was being produced. aPCR step: the biotinylated dsDNA template was then used with a single primer, to generate ssDNA encoding one of the strands used to produce the final EDGE construct. Streptavidin pulldown: the biotinylated dsDNA template was removed via binding to streptavidin, leaving a purified ssDNA product. Hybridisation: to form our triplex construct, we hybridised the ssDNA products for ‘top’ and ‘bottom’ strand. (b) Agarose gel electrophoresis (3%, 120V, 50 minutes run time with ice cooling, GelRed staining). ssDNA components had been extracted from a preceding gel. (c) Fluorescence intensity after 1000 minutes of transcription at pH8 for the original TFR-only (non-triplex forming control), T7D and T7P constructs. (d) Transcription kinetics for the three constructs at six different pHs. For (c) and (d): each data point is the mean of three replicates and the error bar is the standard deviation. Buffer = 1x 100% AceOTB.

We used single-stranded components purified by gel extraction to prepare a construct with no triplex-forming tail (‘TFR only’), as well as a T7P construct (9nt linker, 40nt triplex) and a T7D construct (54nt linker, 40nt triplex, 90% T•AT), and tested transcription in AceOTB of various pH levels, measuring iSpinach fluorescence output for 1000 minutes. Transcription generally proceeds more rapidly at approximately pH8, as expected for the T7 polymerase.

The observed signal is strongly pH-dependent, due both to the innate pH-dependence of transcription speed and the pH-dependence of iSpinach output (Extended Data Figure 1). The effectiveness of the triplex at inhibiting transcription can be measured by examining end-point fluorescence (Fig. 3c) or the kinetics (Fig. 3d). While T7P (9nt linker, 40nt triplex) was unable to inhibit transcription compared to the non-triplex control, the T7D design (54nt linker, 40nt triplex) was capable of inhibiting transcription at all pH levels tested. Furthermore, the T7P approach also provided little customisability of the triplex-forming sequence due to the consensus promoter sequence needed for transcription. Hence, all our subsequent constructs were of the T7D form, with the triplex downstream of the promoter.

The finding that T7D inhibits transcription more than T7P is consistent with our earlier measurements (Fig. 2c), which showed that the T7D triplex remained intact in AceOTB even up to pH8.5, whereas the T7P triplex was evidently less stable. However, the previous measurements suggested that the triplex would be mostly closed in conditions under which we see transcription. We hypothesise that the forces exerted by a polymerase can play a role in opening the triplex, meaning that we see successful transcription under pH conditions that would only partly destabilise a triplex in the absence of the polymerase. This is plausible because calculations suggest that the work done by an RNA polymerase during the scrunching [37] operation associated with transcription initiation is similar to the size of the Gibbs free energy of triplex formation (Δ*G*TFO). To elaborate: using the empirical model of Roberts & Crowthers [38], and excluding the loop contribution, we find that Δ*G*TFO for a representative triplex (90_TFR40_42) at pH7.5 is approximately 1.6×10^-19^J/molecule. We compute this value by noting that there are 36 Ts, 4Cs and 0 CC dinucleotides in this triplex, and substituting into Eqn. 2b in Ref. [38]. In contrast, the work done by a polymerase during scrunching is of the order of 0.4-1.1×10^-19^J/molecule (assuming a force of 10-30pN and a scrunching distance of 11bp i.e. 11×0.34nm=3.74nm, and applying work = force × distance gives 37-112pN nm i.e. 37-112×10^-21^J). Further credence is given to our hypothesis by the experimental proof that protein machines can unravel complex nucleic acid structures, with one study even demonstrating that a ribosome could forcibly dismantle an RNA-DNA origami and perform translation from an mRNA sequence that had been packaged within [33].

We can eliminate the possibility of melting due to temperature rises associated with dissipation of electrical power because the heat released could only raise the temperature by a fraction of a degree on average. We calculate this as follows. The current *I* is kept constant at 130μA and the voltage *V* is approximately 3V, giving power *P=IV=*130μA×3V=0.39mW. The power is maintained for a time, *t*, of 107 seconds at most. Hence, the energy, *U*, is no greater than *U = Pt =* 0.39mW×107s = 42mJ. We may assume that the transcription mixture has the same heat capacity, *C*, and density, *ρ*, as water (*C*=4.2 J g^-1^K^-1^, *ρ=*1 g mL^-1^), and the reaction volume, *V*, is 50μL. The heat transferred is Δ*Q* = *CρVΔT,* where *ΔT* is the temperature change we wish to calculate. Assuming that all electrical energy is converted to heat, rearranging the equation for Δ*Q* and substituting in the appropriate values, we obtain a temperature rise of 0.2°C.

### Hardware production and calibration

We generated a library of EDGE constructs (Fig. 4a), with all 48 possible combinations of four triplex lengths (10nt, 20nt, 30nt and 40nt), four base compositions (60%, 70%, 80% and 90% T•AT), and three linker lengths (42nt, 48nt and 54nt). These were assembled in an automated facility (the Edinburgh Genome Foundry), together with the corresponding non-triplex-forming control constructs. Due to the increase in scale and 96-well plate format, the protocol was altered to be more automation-friendly, switching to vacuum pump purification instead of centrifugation and using fluorescent dye-based quantification instead of UV/vis spectrophotometry. The gel extraction step was also omitted.

**Figure 4:**
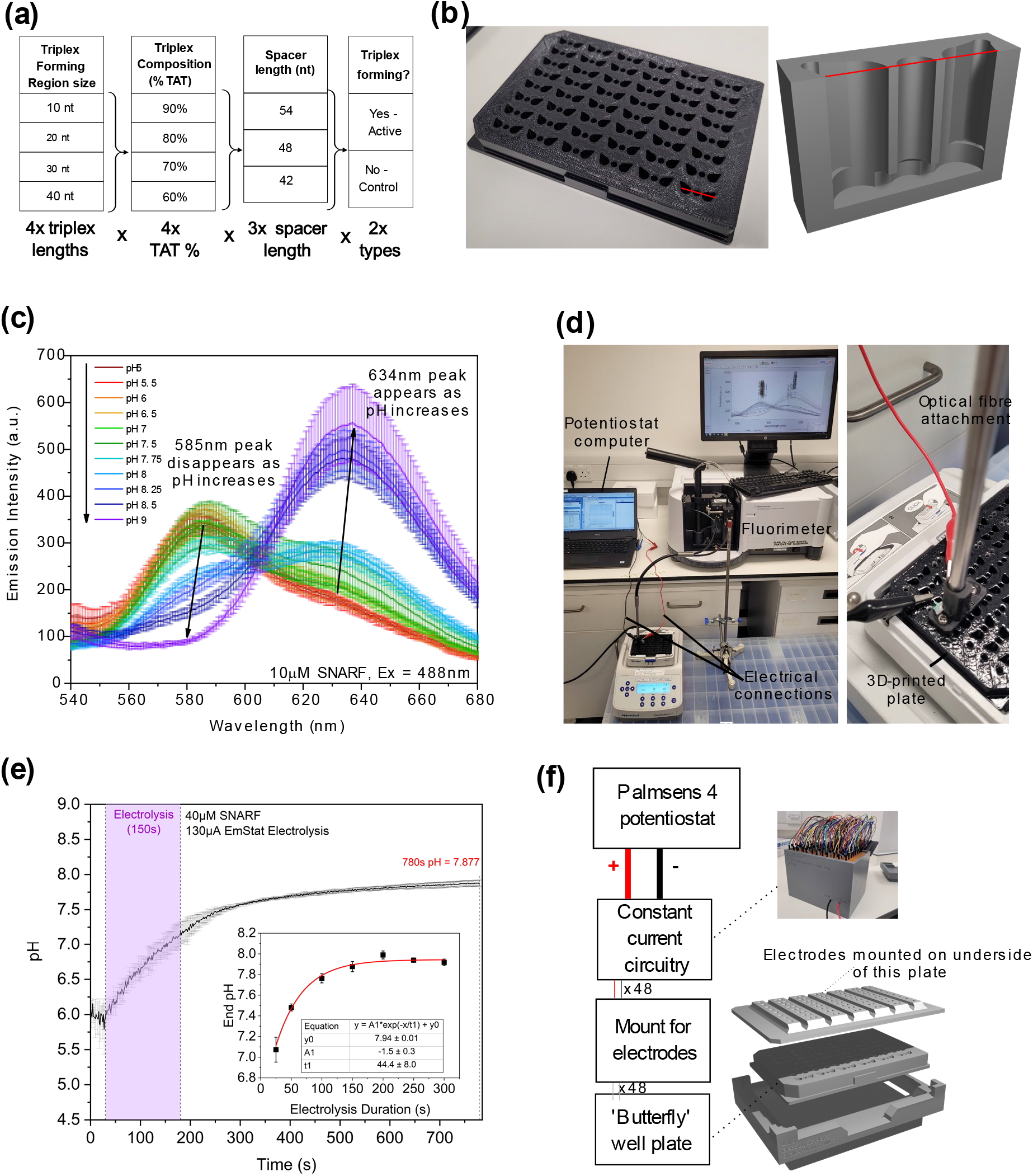
(a) The combinations of parameters used for assembly of constructs. (b) Photograph of acetone-smoothed 3D-printed ABS plate used for high-throughput experiments and rendering of 3D cross-section. (c) Representative measurement of fluorescence of SNARF dye under excitation with light at 488nm at various pHs, showing the disappearance of the 585nm emission peak and the appearance of the 634nm emission peak as pH increases. Measurements were performed in quadruplicate in AceOTB at different pHs. Error bars: standard deviation. (d) Setup in which the fibre optic attachment of the Agilent Cary Eclipse fluorimeter was used to measure the fluorescence of the SNARF dye during electrolysis, driven by a PalmSens 4 potentiostat. (e) pH before, during and after electrolysis of AceOTB, during a representative run, measured using SNARF fluorescence, where the 585nm/634nm emission ratio was calculated as a function of pH and fitted using a sigmoidal function. Seven different durations of electrolysis were tested, giving ‘end pH’ (10 minutes after end of electrolysis) as shown in the inset. Error bars = standard deviation (*N*=3). Buffer = 1X 1% AceOTB. (f) Setup used for high-throughput characterisation of EDGE constructs. The grey box in the photograph contains the constant current circuitry, and the wires emerging from it are connected to pairs of electrodes in a black plate (just visible behind the box). The electrode plate is then lowered into a butterfly-well plate of the form shown in part (b) and in the exploded diagram.

To test electrically controlled transcription from these constructs in a high-throughput manner, we required bespoke hardware that was capable of electrolytic pH shifting while also being compatible with automated liquid handlers and plate readers for high-throughput testing. We employed fused deposition modelling (FDM) 3D printing to prepare custom 96 well plates that connected pairs of wells via an integrated salt bridge (Fig. 4b). Plates were printed using black acrylonitrile butadiene styrene (ABS) filament, as ABS can be acetone smoothed, where the outer ABS surface forms a slurry with acetone vapour and ‘melts’ [39]. This allowed the layer lines produced during 3D printing to be smoothed to produce a more reliably watertight part for use with liquid reagents. ABS is also acid resistant, allowing the plates to be acid cleaned prior to use with *in vitro* transcription reactions.

The design of the wells was an angled teardrop, which allowed for electrodes to be inserted into the solution without entering the light path during fluorescence scanning. The customised nature of the plates also allowed for the size of the wells to be designed for the intended reaction size (50µL + 30µL mineral to prevent evaporation), and included channels that connected the floors of wells in pairs. These channels were filled with molten agarose via a central opening that then cooled and set to form a salt bridge that limited diffusion between the wells while completing the circuit. By having the salt bridge contained within the plate, the electrolysis procedure was simplified, limiting the number of parts to be introduced into the wells from above. 3D printing is the only method that enables this plate to be prepared as a single piece due to the overhanging structures involved in forming the channels inside the part.

We needed to develop an electrical system that would be capable of bringing the transcription solution to a precisely determined pH. For such calibration purposes, we used the fluorescent dye SNARF (excitation wavelength 488nm), which emits at 634nm at high pH and 585nm for low pH (Fig. 4c). These measurements were taken on SNARF in different pH AceOTB solutions, within a variant of our custom plates, using the fibre optic attachment for our fluorimeter (Fig. 4d). The data was used to provide a calibration curve, enabling us to determine pH of an unknown solution based on the emission spectrum of SNARF. By measuring SNARF emission spectra as a function of time, we were able to establish how pH changes after each electrolysis (Fig. 4e). We ascertained how the final pH depends on the length of electrolysis at a constant current of 130μA (Fig. 4e, inset), which enabled us to estimate the length of pulse necessary for subsequent experiments – 54 seconds to move from pH6 to 7.5. We constructed constant-current circuitry (Fig. 4f and Supplementary Information) to deliver an electrolysis pulse of a set duration at a current of 130μA. For the large-scale experiments with the EDGE constructs, we ultimately started at pH6.6 rather than pH6 and a 54-second constant-current 130 μA pulse took the pH to 7.8.

### Response of different design variants

We measured iSpinach fluorescence kinetics for two hours for cell-free transcription reactions involving either an EDGE construct or the corresponding control structure. In each reaction, we set the pH using electrolysis, with a current pulse of either 0s, 26s, 54s or 107s in duration. For every construct and every electrolysis duration we performed the experiment in triplicate, before averaging and fitting the data (Fig. 5a, Methods). The output of our analysis procedure comprised a set of 384 amplitudes, corresponding to 4 electrolysis durations for each of 48 triplex constructs and the same for each of 48 control constructs.

**Figure 5:**
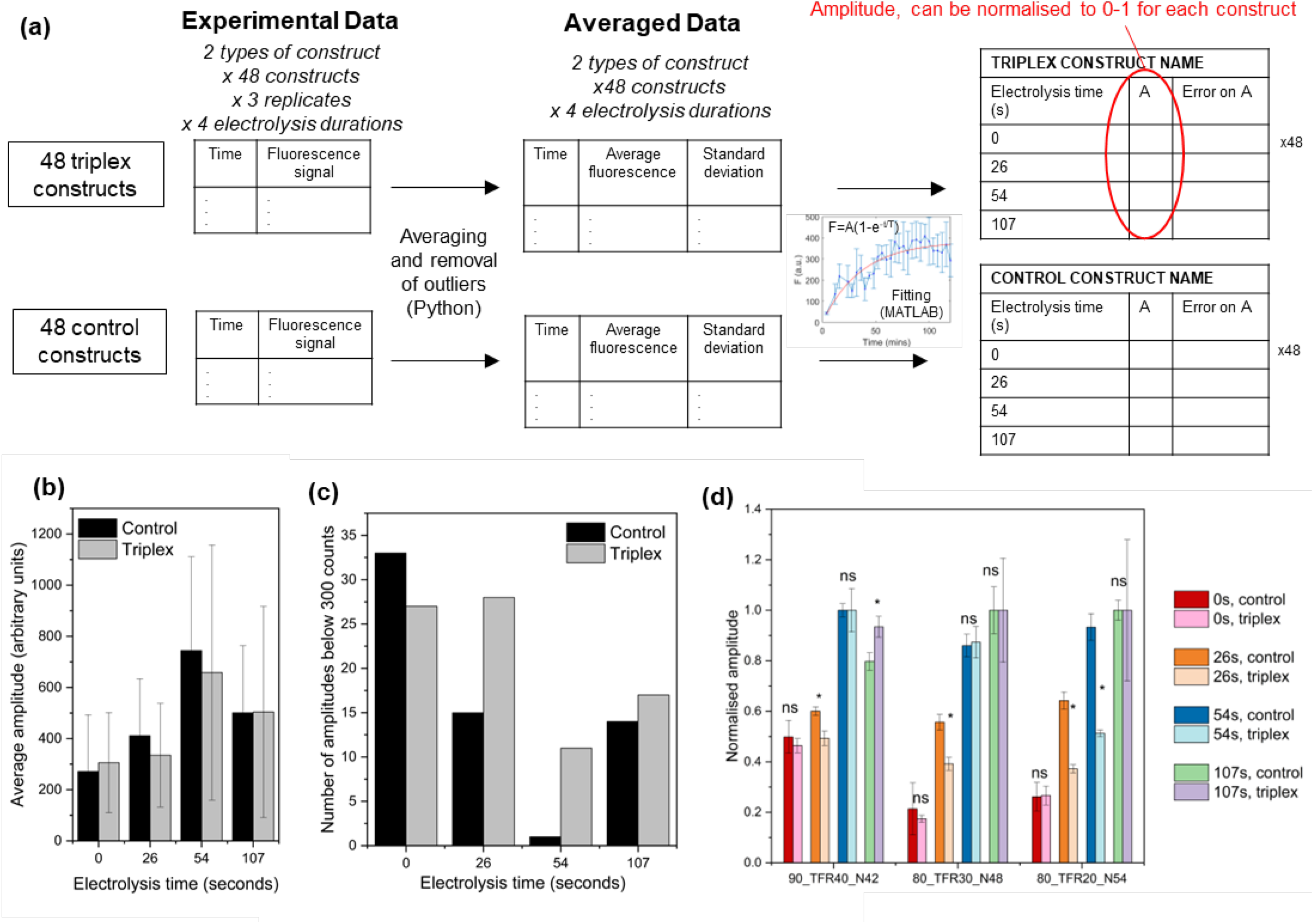
(a) Schematic diagram showing data analysis procedure. For each construct and each electrolysis condition, data for each set of replicates was averaged to give a kinetic profile that was then fitted using MATLAB, yielding fit parameters that were then analysed. The inset graph illustrates a representative dataset, the fit function and the result of the fit. The amplitude parameter is indicative of the level of transcription observed from the construct. (b) Average (mean) amplitude across all 48 constructs of each type, for each electrolysis time. The error bar represents the standard deviation. (c) The number of cases in which the amplitude (from the fit) was below 300 counts (arbitrary units). (d) Normalised amplitudes for three scenarios of interest; for each construct, we normalised the set of four amplitudes corresponding to the different electrolysis durations, yielding values on a 0-1 scale. Statistical tests then confirmed where the control and triplex constructs exhibited different behaviour. Working and p-values are given in a supplementary file.

In our experiments, the measured amplitude could have been affected by numerous factors other than the state of the triplex unit. For instance, while we aimed to use the same concentration of DNA construct in each reaction, the methods used to mass-produce constructs did not achieve the same level of purity as we would have achieved in one-off preparation of a single construct, meaning that quantitation was not as accurate. Furthermore, the need to perform measurements on a large number of constructs simultaneously in a practical length of time placed constraints on the design of our hardware that resulted in more variability than would be seen in a typical plate measurement. We thus see a trade-off between quantity and quality of data. If we had worked with a smaller number of constructs, it would have been possible to achieve a higher level of purity and superior signal-to-noise, but we would have been sampling a smaller design space. With no *a priori* information regarding what length, T•AT content and loop length would be required to achieve the transcription-modulation effect we needed, it was necessary to sample as large a design space as possible, which required us to test as many constructs as possible.

The amplitude observed in our experiments could also have been affected by any sensitivity to the sequence flanking the promoter, as well as the intrinsic pH-dependence of transcription. As described earlier, we isolate the effect of the triplexes by comparing the performance of each triplex construct with that of a control construct that shares exactly the same sequence apart from in the tail. In the case of the control construct, the tail is incapable of forming a triplex. Both the control construct and triplex construct should be affected in the same way by the sequence surrounding the promoter and the intrinsic pH-dependence of transcription, meaning that differences between them are attributable to the effect of the triplex. By drawing on such a comparison, we are also able to avoid the compounding effect of the pH-dependence of DFHBI fluorescence (Extended Data Figure 1). While DFHBI yields a substantial signal at all the pH levels to be reached in our experiments, the signal does decrease at lower pHs, which could complicate analysis in the absence of controls.

It is instructive to consider the average amplitude across all the control constructs for each of the electrolysis durations tested (Fig. 5b). For each duration, we found considerable variation (large standard deviation) across the set of all constructs. This is to be expected due to the factors mentioned above i.e. hardware variability, quantitation inaccuracy and effects related to changes in the sequence flanking the promoter. Given the size of the standard deviation, there can be more variation between one control construct and another at the same electrolysis point than between the same construct at different electrolysis points. However, we saw that the average amplitude was generally lowest when there has been no electrolysis, as the pH is sub-optimal for transcription. Increasing the electrolysis duration to 26s increased the signal and a further increase was seen for electrolysis duration of 54s. This is reasonable because these electrolysis conditions have increased the pH such that conditions are optimal for transcription, while also increasing the signal for DFHBI. When the electrolysis duration is extended to 107s, the average amplitude drops, which we attribute to the pH being too high for optimal transcription. Given the variability of amplitudes, there is no further insight that can be gained from averages taken across the whole design space.

While caution is required in the interpretation of absolute values of amplitudes, due to issues such as the quantitation inaccuracy noted above, we may use our data to ascertain whether there are any cases in which the triplex appears to be completely suppressing transcription under conditions that should be optimal. We therefore considered the number of amplitudes observed to lie below 300 counts across the whole data set (Fig. 5c). We chose the level of 300 counts because this is close to the average baseline for zero electrolysis. We found that the number of amplitudes below 300 counts was similar for control and triplex constructs at both 0s and 107s of electrolysis. However, there is a marked difference at 26s and 54s. Here, we found that far more of the triplex constructs yield low signals. For 54s of electrolysis, only 1 control construct had an amplitude below 300 whereas 11 triplex constructs did. This suggests that as many as 11 of our triplexes may have been acting as brakes on transcription at this pH. Using the notation % T•AT_TFR(triplex forming length in nucleotides)_N(loop length), the constructs in question were: 90_TFR20_N48, 80_TFR40_N54, 80_TFR40_N48, 80_TFR40_N42, 80_TFR30_N54, 80_TFR10_N54, 70_TFR40_N54, 70_TFR40_N48, 70_TFR20_N48, 60_TFR40_N42, 60_TFR10_N54. Interestingly, these constructs spanned much of the design space rather than being clustered in specific regions. We note that the average length of triplex in these constructs was 30nt.

We also found that some of our triplex constructs were not suppressing transcription in any meaningful way. In our datasets, a signal of 900 can be regarded as ‘high’, and after 54s of electrolysis 11 triplex constructs yielded a final amplitude of over 900 (90_TFR20_N54, 90_TFR20_N42, 90_TFR10_N54, 90_TFR10_N48, 90_TFR10_N42, 80_TFR30_N48, 80_TFR20_N48, 80_TFR20_N42, 80_TFR10_N48, 60_TFR30_N42, 60_TFR20_N42). The average length of triplex in these constructs was approximately 18. Consequently, it appears that longer triplexes are more likely to be effective brakes on transcription, which is reasonable as we would expect a longer triplex to be more stable and less likely to open for transcription. We also note that some of these constructs differ from some members of the previous (fully suppressed) group in only loop length (e.g. 80_TFR30_N48 vs 80_TFR30_N54), suggesting that relatively small changes in design can dramatically change the observed behaviour. There is also clearly an interplay between the design factors because 90_TFR20_N48 is inhibited and 90_TFR20_N54 is active, but in the case of 80_TFR30, we found that the 48nt loop construct was active and the 54nt loop construct was inhibited.

The two foregoing groups of constructs did not achieve our goal of actively modulating transcription under conditions we can access. To identify constructs that do this, we needed to adopt appropriate screening criteria. We sought triplex constructs that yielded some transcription after 54s of electrolysis, but where the triplex amplitude for 54s and/or 26s was suppressed somewhat in comparison with the corresponding control. In order to compare electrolysis-induced changes seen for the triplex amplitude with those of the corresponding control, normalisation is appropriate. In this way, for each triplex we examined a set of four amplitudes (for each of the four electrolysis durations) on a scale of 0-1 and compared them to the corresponding set of four amplitudes lying on a 0-1 scale for the control. By normalising the amplitudes, we avoided variation arising from concentration differences.

Examples of cases meeting our criteria are shown in Fig. 5d. Here, we have performed a 2-tailed t-test to confirm which differences were significant (based on a *p*-value threshold of 0.05, following convention). The clearest example of the desired behaviour is displayed by 80_TFR20_54. Interestingly, both of the other two 80_TFR20 constructs gave high signals after 54s of electrolysis. This suggested that for a 20-nt triplex with 80% T•AT content, the tension in the 42nt and 48nt linkers was too great to sustain a stable triplex. In contrast, when the T•AT content is 90%, the triplex was stronger, meaning that a 48nt linker could support a 20nt triplex and lead to a construct where transcription is fully suppressed.

## Conclusion

Overall, we conclude that a pH-sensitive triplex can act as an effective brake on transcription, which enables us to perform ‘Electrically Directed Gene Expression’, where transcription is activated or deactivated by electrolytic changes in the reaction medium. EDGE is a platform technology that could be used wherever there is a need to modulate RNA production rate in a cell-free system. For example, we may conceive an information processing system that receives electrical inputs (in this case an electrolysis pulse) and converts them into biomolecular triggers (here, an RNA signal) that feed into nucleic acid logic gates based on strand displacement. We have also considered the application of EDGE in the production of mRNA vaccines, as a method for modulating transcription. A third application might involve a biomaterial undergoing a change of state in response to the generation of an RNA strand, produced by transcription activated by an electrolysis-induced pH change. However, the applications may be limited by the requirement for the buffer to be relatively weak as well as the need to induce a significant pH shift to affect transcription output.

.The methods we developed to demonstrate EDGE may have broad applications, as our work illustrates both the use of 3D-printing to create customised laboratory plasticware as well as a method for interfacing electronics with cell-free systems in a high-throughput experiment. Our plasticware could not be constructed using injection moulding, as layer-by-layer bottom-up manufacture is needed to provide the required internal structure for the agarose bridge between the wells. Our early experiments with stereolithography as a 3D-printing modality revealed two critical issues – the fluorescence of the calibration dye SNARF was distorted by the resin and the structure was also not watertight. When we used fused deposition modelling (FDM) with ABS filament, we were able to deploy acetone smoothing to render the plate watertight and avoid leakage. FDM is also a cheaper production method and the high chemical resistance of ABS enabled us to use acid cleaning to sterilise the wells before use. Our production method inevitably had more variability than scaled-up commercial manufacturing processes, with acetone smoothing and FDM layer lines introducing well-to-well variability. In future work, these effects could potentially be reduced further by optimisation of the plate design and adjustments to the transcription mixture recipe to increase the observed signal. Despite the comparatively high level of variation and noise, our work serves to illustrate the utility of 3D printing for prototyping and proof-of-concept demonstrators in a biochemical context, mirroring the known uses for bespoke manufacturing in settings such as healthcare [40]. More evolved versions of our platform would most likely be based on miniaturised architectures, comprising a lab-on-chip system with microfluidics for liquid handling and microelectrodes for control of pH.

Finally, we speculate that it might be possible to develop our EDGE system further to enable electrical control over translation as well, although this would be significantly more complicated and is beyond the scope of the present work. If cell-free protein synthesis could be controlled electrically, it could be possible to design implants that would produce therapeutic proteins such as cytokines or enzymes on demand in response to electrical stimuli.

Future work could explore the use of more advanced computational approaches to guide the design of DNA-based switches for transcriptional regulation, building on our use of NUPACK [41] and EGNAS [42], as well as the achievements of others in the implementation of computer-aided DNA strand design in a variety of contexts [43, 44, 45, 46] Other priorities for future work could include the development of new miniaturised lab-on-a-chip hardware to control the samples, as described above, as well as extending the methodology to enable repeated electrolysis-driven switching of transcription; our existing attempts to achieve this (data not shown) suggest that one or more of the components degrades under the conditions used.

---

**Extended Data Figure 1:**
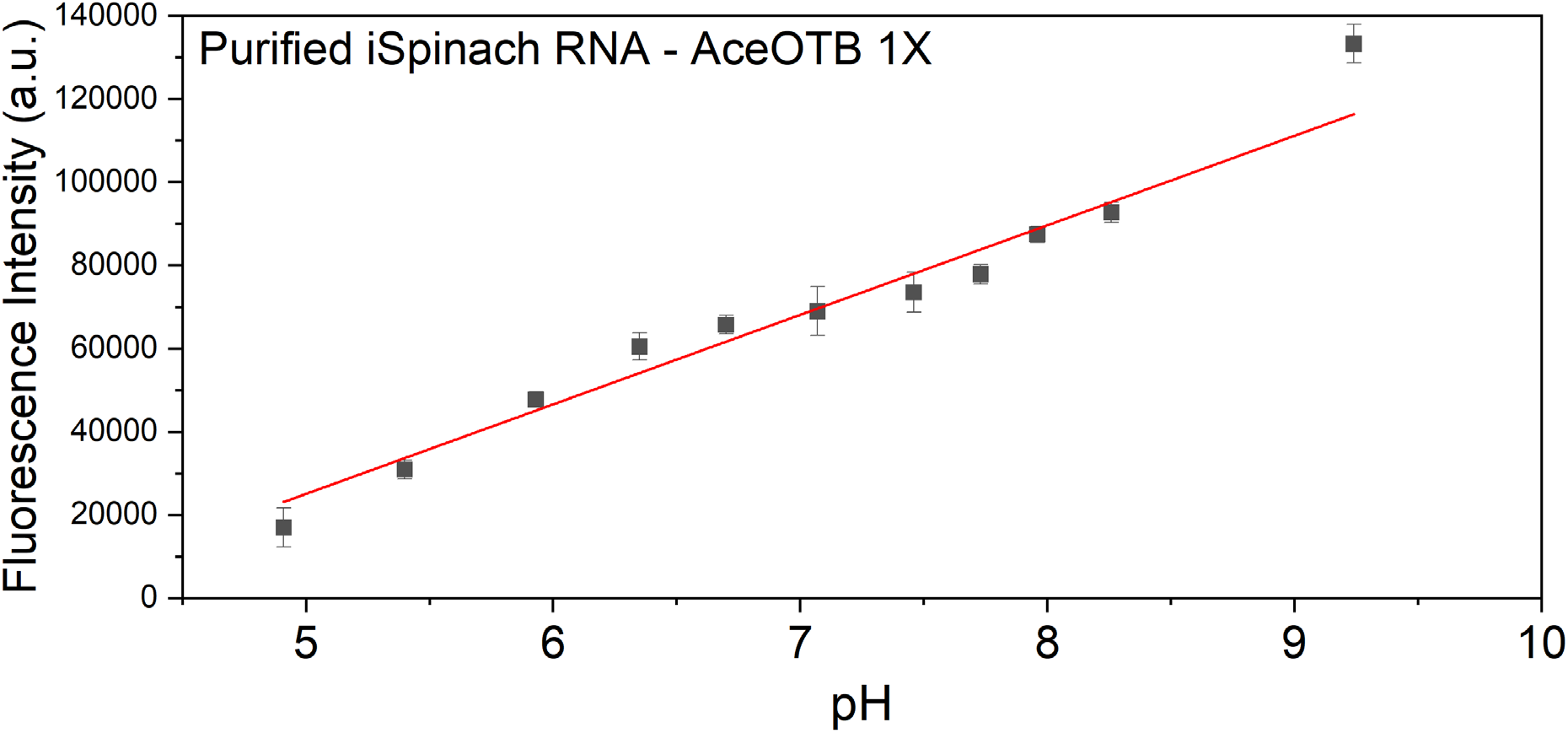
Fluorescence of purified iSpinach/DFHBI in different pH AceOTB buffers. iSpinach has strong fluorescence across the pH ranges used in this work, but it does demonstrate pH dependency that will affect its fluorescent output, potentially exaggerating the apparent pH-dependence of transcription effectiveness. Data was measured in triplicate and the error bars represent the standard deviation.

**Extended Data Figure 2:**
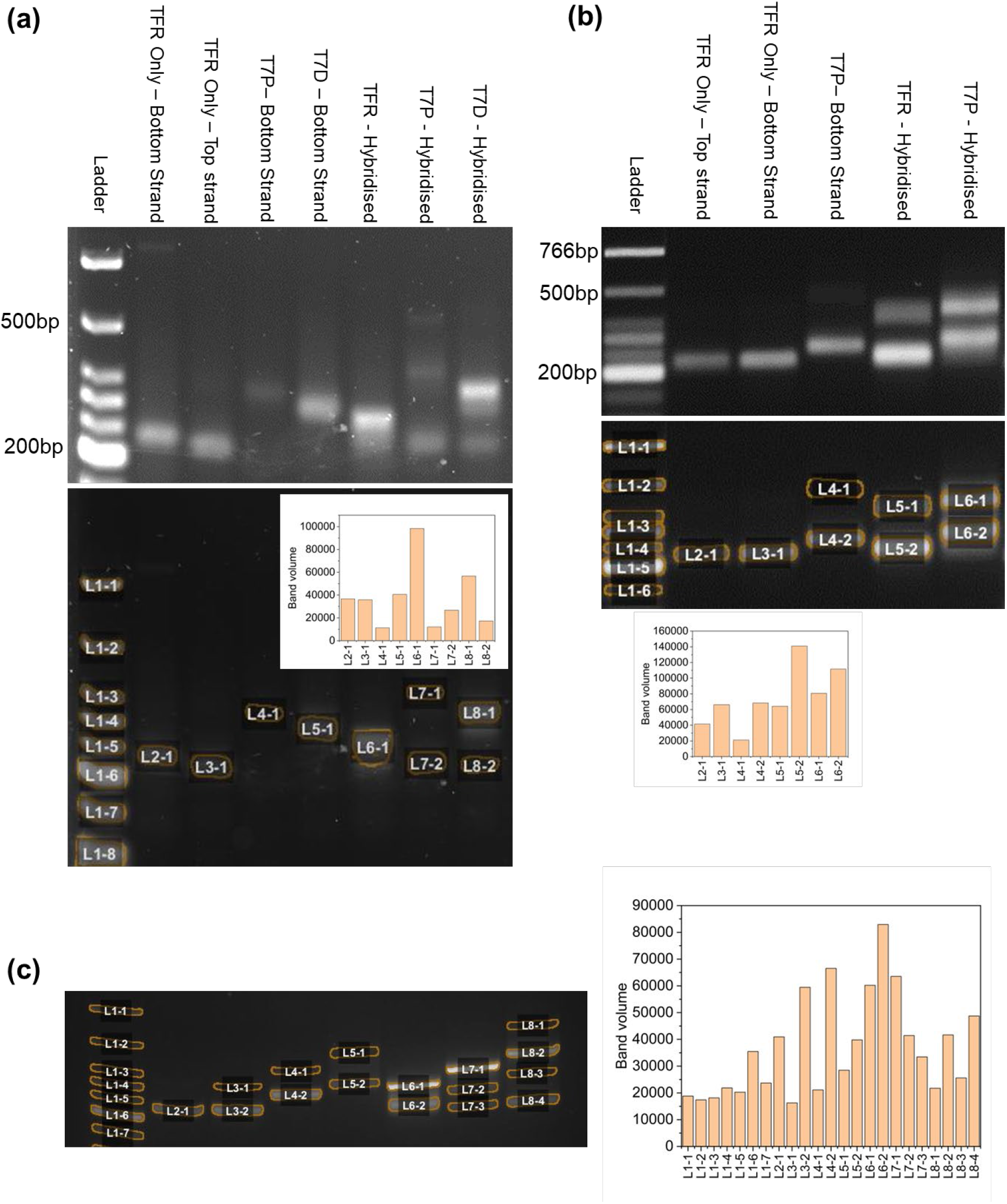
in-depth analysis of gel electrophoresis of selected constructs, using gel images recorded during optimisation of experimental conditions. Analysis was performed using open-access AI-powered tool GelGenie v1.0.5.1 with QuPath 0.5.1 (https://github.com/mattaq31/GelGenie & [47]), with the default settings. Bands are detected by segmentation. The top image in each panel shows the gel without any band labels, while the bottom image shows the bands identified by GelGenie’s segmentation and the graph shows the intensity of each band as quantified by the software. In most cases, lanes containing a single strand have one well-defined band, as expected. On occasion a second band appears, as in (b), T7P-bottom strand (L4-1). These bands are presumably the result of secondary structure formation or association of individual strands into multi-strand complexes. Lanes containing hybridisation products generally contain two well-defined bands, one associated with the intended construct and the other associated with the excess of one of the inputs. Different samples/gels show some variation in terms of band intensity and the presence of off-target complexes/structures due to minor unavoidable differences in concentration of DNA. (a) Gel acquired during experiments to optimise the number of PCR cycles; 12 cycles were used in this instance (later runs utilised 14 cycles). 3% gel, run at 75V for 1h, with 30 min post-run stain using GelRed (b) Gel acquired during optimisation of large-scale production, for testing of purification using Qiagen^TM^ vacuum filtration with QIAquick^TM^ 96-well plates. 3% gel, run at 90V for 1 hour, GelRed added to samples prior to run. (c) Application of GelGenie processing to the gel provided in Fig. 3b.

## Materials and methods

DNA sequences, data, code, and details of electrical circuit design are provided in the supplementary information. Further information is available in Ref. [48].

### Triplex sequence design

Triplex sequences (both TFR and TBR) were designed to form a triplex that favoured only a single configuration in order to ensure triplex of the intended length formed, with a unique pattern of T•AT triplets between each C+•GC triplet. Linker sequences were generated using the Exhaustive Generation of Nucleic Acid Sequences (EGNAS) tool (Version #1158) to ensure that the full tail length did not form a secondary structure with the TFR and remained fully single-stranded [42]. EGNAS was also used to generate non-triplex sequences to replace the TFR in control constructs with sequences that would not form secondary structures to the previously designed linker regions. Generated sequences were checked using the Nucleic Acid Package [49] (NUPACK) online tool to confirm the lack of secondary structures within the ssDNA tail.

### Fluorophore investigations (DNA sequences provided in Supplementary Information section 1.1.1)

Two oligonucleotides were synthesised by Integrated DNA Technologies that once hybridised could form a 10nt triplex, with a Cy5 fluorophore at the 3’ end of the ssDNA TFR and a Black Hole Quencher 2 (BHQ2) quencher at the 3’ end of the dsDNA TBR. This was placed into a cuvette of 1M sodium sulphate with a final concentration of 25nM of hybridised construct. Using a multi-cuvette adapter a second cuvette of 1M sodium sulphate was connected via a salt bridge made of filter paper soaked in 1M sodium sulphate, with a graphite electrode lowered into each cuvette from above using a 3D-printed mount. The mount also included a small motor connected to a polystyrene stirrer that lowered into the DNA cuvette to speed up diffusion. Fluorescence kinetics were scanned using an Agilent Cary Eclipse fluorescence spectrophotometer (excitation = 648nm, emission = 667nm), and electrolysis was performed using the voltage source unit of a HPTM4156B semiconductor parameter analyser. A voltage of 5V was applied for 3 minutes, before reversing the polarity and electrolysing again using the same voltage and duration.

T7P and T7D fluorophore oligonucleotides were designed and tested in a similar manner, using Britton Robinson buffer prepared to known pH levels instead of sodium sulphate and without electrolysis. The T7D construct was assembled out of 3 hybridised oligos instead of 2, due to length limitations on fluorophore labelled oligo synthesis.

### PCR / aPCR

DNA sections to be amplified were synthesised as either IDT linear DNA gBlocks or as Invitrogen GeneArt plasmids (sequences provided in Supplementary Information file section 1.1.2). Primers were designed using either the IDT PrimerQuest online tool or a self-produced primer design Python script (provided in Supplementary file, sequences given in Supplementary Information section 1.1.3). If being used to produce biotinylated DNA template for aPCR, one of the primers would be biotinylated. This would be the reverse primer if producing top strands, and forward primer if producing bottom/tail strands. PCR and aPCR reactions were performed using Q5 polymerase (NEB - #M0491L), starting with 4 x 10^-13^ moles of dsDNA template and including GC enhancer (NEB - #B9028A). PCR reactions were then purified via Qiagen QIAquick PCR purification columns, using spin columns if performed by hand (Qiagen - #28106) using a centrifuge (Hitachi - #CT15RE) or using AMPure magnetic DNA purification beads (Beckman Coulter - #A63882) if performed by the Edinburgh Genome Foundry (EGF). Before purification, aPCR reaction solutions were loaded into Pierce Streptavidin plates (Thermo Scientific - #15500) that had been previously washed according to the manufacturer’s protocol and left shaking for 4 hours at 400rpm. aPCR reactions were then purified via Qiagen QIAquick PCR purification columns, using spin columns if performed by hand (Qiagen - #28106) or in a 96-well plate vacuum column format if performed by the EGF (Qiagen - #28181). Additionally, ssDNA used during the small-scale chemical control tests were not purified via streptavidin pulldown, but were separated via agarose gel electrophoresis, excised and purified using a gel extraction kit (NEB - #T1020S). After aPCR, ssDNA products were quantified using QUBIT dye and fluorescence spectrophotometry using either an Agilent Cary Eclipse fluorescence spectrophotometer if using cuvettes or a BMG Clariostar if using plates (excitation = 485nm, emission = 527nm). As the interactions of QUBIT dye with the secondary structure of each ssDNA were unknown, each ssDNA product had an individualised ratio of fluorescence to a known quantity of ssDNA calculated. UV-visible spectrophotometric quantification of Qiagen QIAquick-purified ssDNA was used as a reference for the fluorescence value of the Qubit dye, using NUPACK simulations of each potential aPCR product to generate a custom extinction coefficient to improve UV-vis quantification accuracy (Supplementary Information section 1.1.4). Top and bottom strands were standardised to 50nM and hybridised together by heating to 95°C in a thermal cycler and cooling to room temperature.

### AceOTB preparation

AceOTB is based on a previously published optimised transcription buffer [50]. The alteration to ‘AceOTB’ is the substitution of chloride ions for acetate in order to avoid producing chlorine gas while electrolysing. There are two versions of AceOTB; ‘100%’ and ‘1%’. This refers to the amount of tris base, with ‘100%’ being the standard recipe as per the original reference. ‘1%’ refers to using ‘1%’ of the original amount of tris base to make a much more weakly buffered solution that requires less electrolysis to shift its pH level. Both varieties of AceOTB were produced at 10X concentration for dilution during reaction assembly. 10mL of 10X (100%) AceOTB contains 500mg Trizma (Sigma-Aldrich - #T1503-25G), 128.5mg Magnesium Acetate (Fisher Scientific - #423875000), 16mg DTT (Fisher Scientific - #327190010), and 3mg Spermidine (Alfa Aesar - #A19096.03). These are suspended in 5mL ultrapure water, and adjusted to the desired pH using 3M acetic acid (Fisher Scientific - #423225000) before filling the remaining volume up to 10mL with ultrapure water. For 1X (100%) AceOTB, the 10X solution was diluted ten-fold. For the 10X 1% AceOTB, the quantity of Tris base was reduced to 1/100 of that given in the 100% recipe and pH was adjusted by titration with acetic acid in the same way.

### In vitro transcription

Transcription solution was prepared in bulk as follows (per 50 µL transcription reaction): 5µL AceOTB 10X, 2.5µL rNTPs (NEB - #N0466S), 1µL T7 RNA polymerase (NEB - #M0251S), 1µL 2mM 3,5-difluoro-4-hydroxybenzylidene imidazolinone (DFHBI, Sigma-Aldrich - #SML1627-5MG) (DFHBI is suspended in DMSO (Sigma-Aldrich - #D8518-50ML)), and 30.5µL ultrapure water. This was set up in triplicate across 3 custom EDGE plates using a Beckman Coulter Biomek FXP liquid handler, with 10 µL of 25nM hybridised EDGE construct DNA, followed by 30µL of mineral oil (Sigma-Aldrich - #M5904-500ML) added to each reaction well. iSpinach fluorescence was scanned using a BMG Fluorostar Omega (Excitation = 485nm, emission = 520nm), with a PAA SCARA KiNEDx robotic arm shuffling which plate is being scanned throughout the reaction, allowing kinetics to be scanned across three plates simultaneously.

### RNA purification

iSpinach RNA was transcribed in a bulk 1000µL reaction mixture and purified using a ReliaprepTM RNA purification kit (Promega - #Z6010). The RNA was then added in equal quantities to AceOTB with DFHBI at various pH levels. This was heated to 95°C and allowed to cool to room temperature to ensure correct folding of the iSpinach structure before scanning for fluorescence in a BMGTM Clariostar (excitation = 480nm, emission = 520nm).

### Electrolysis calibration

The strength and duration of electrolysis was calibrated using a Palmsens 4 potentiostat. A voltage of 3V was chosen as a compromise between speed of pH shift and potential detrimental effects of high voltages on the components of the *in vitro* transcription solution. Chronoamperometry determined that in the standardised 3D-printed setup this was achieved with a current of ∼130μA across each well, and that constant current application (rather than constant voltage) produced more reliable pH shifting, taking into account salt bridge and resistance variability between pairs of wells. The pH of the solution was calculated using 5-(and-6)-carboxy SNARF-1 (Thermo Fisher ScientificTM - # C1270) fluorescent dye that shifts its emission peak depending on pH. This was measured using an Agilent Cary Eclipse fluorescence spectrophotometer to measure both peaks (excitation = 488nm, ‘low pH’ emission = 585nm, ‘high pH’ emission = 634nm) while being electrolysed, using the ratio between the emission peaks to calculate the pH over time. To electrolyse while scanning fluorescence in the same setup to be used in large scale experiments, a fibre optic probe attachment for the Agilent Cary Eclipse fluorescence spectrophotometer was installed and fitted to an additional 3D-printed fibre optic connector variant (FOCV) design of EDGE plates that replicated the reaction setup but included screw holes for mounting the fibre optic probe (with an additional SLA printed adapter), along with a pair of platinum electrodes (FIG) inserted from above. This setup was used with the chronopotentiometry mode of the Palmsens 4 potentiostat to produce a calibration curve of duration electrolysis and induced pH shift using a constant current of 130µA. Further details of the optimisation of the electrolysis process are given in Chapter 5 of Ref. [48].

### Custom hardware design and production

Parts were designed using Autodesk Inventor 2019 (v2019.1.1) and 2022 (v2022.0.1). EDGE plates were sliced into GCODE using Raise3D Ideamaker software (v4.2.2) with print settings of a nozzle diameter of 0.4mm, layer height of 0.15mm, 10\% gyroid infill, nozzle temperature of 250°C and bed temperature of 100°C. The print layout also included a raft underneath each plate to help with bed adhesion due to the tendency of unsecured ABS parts to contract and curl, pulling themselves off the printbed. Plates were then printed on a Raise3D E2 printer using 1.75mm Black ABS filament (RS PRO - #832-0311) in duplication mode, allowing two printheads to mirror each other and print two plates at a time. Black was chosen to help reduce background and crosstalk.

After printing, plates were acetone smoothed using a plastic box with a 40mm 12V fan attached to the underside of the lid. Plates were placed 3 at a time in the box using a 3D-printed PLA rack along with 100mL of acetone split across 2 100mL beakers and left for 90 minutes with the fan running. Plates were then removed and left overnight to air, with excess acetone to be reused another day. The next day, the wells of the smoothed plates were acid cleaned using 2M HCl (ACROS Organics - #10794821) and left for 15 minutes before removing the acid and thoroughly rinsing for 3 times with ultrapure water. Plates were dried overnight in a heated glassware drying cupboard. Agarose salt bridges were made by preparing 0.1M sodium sulphate 1% agarose melted using a stirring hot plate set to 160°C and 350 rpm for approximately 20 minutes until bubbling. A Thermomixer C (Eppendorf - #5382000031) with a plate format SmartBlock (Eppendorf - #5306000006) was loaded with an EDGE plate and preheated to 95°C to warm the plate and prevent premature solidification of the agarose. To each pair of wells, 130 μL of molten agarose was pipetted into the agarose filling channel, with the tip of the pipette not pushed fully into the hole, with the agarose flowing down and not forced in under pressure. Once all channels were filled, the plate was removed from the ThermomixerTM, placed on a flat surface and tapped gently yet firmly to ensure that all of the agarose had flowed through the wells and was level. This was left at room temperature to cool and solidify before being used for EDGE reactions.

The top electrolysis plate was SLA printed. 3D files were sliced into using Chitubox software (v1.6.5.1) with a layer height of 0.025mm, exposure time of 11s per layer, with 8 bottom layers with 160s exposure time, with 100% solid infill and angled to use only bed mounted supports. Parts were printed on an Elegoo Mars SLA printer in black resin (eSUN - #esun-STRES-BLK-1ltr), and washed using isopropyl alcohol (Hexeal - #IPAISOPROPYLALCOHOL). Parts were then cured using an LED UV-lamp or left in direct sunlight. Platinum electrodes were made by cutting platinum wire (Sigma-Aldrich - #349402-1.5G) into 15mm segments, with 5mm being inserted into female end of 20cm male-to-female Dupont connector wires (Avartek - #ACC-CBL-004 M-F) and soldered into place. These were then inserted into the square slots of the printed top plate and secured with a small amount of super glue, leaving 9mm of electrode exposed on the underside of the plate. The electrolysis circuitry (Supplementary Information section 2) used stripboard held within a 3D-printed casing to connect each pair of electrodes to a constant current chip (Texas Instruments^TM^ - #LM334Z) and a 511Ω resistor (Vishay^TM^ - #683-3815) used to calibrate the output current to 130µA per well according to the manufacturer information. All well/chip circuits were connected together in parallel to allow all wells to be electrolysed at once, being powered and controlled by a Palmsens^TM^ 4 potentiostat in chronoamperometry mode. Voltage was set to 5V, however due to each pair of wells being limited individually to 130µA each well was only exposed to approximately 3V.

### Data processing (see also Supplementary Information section 3)

Kinetics data was compiled from the individual Overlord3 output files via Python script (provided in Supplementary Information) to combine replicates for each EDGE construct across plates and calculate standard deviations. This code also removed values that had a standard deviation greater than 50% of the average fluorescence as outliers. The resulting data is provided as a supplementary file (‘EDGE Kinetics Master Sheet.xlsx’), where #DIV/0! represents either a removed outlier point or the baseline time point. The two hour kinetics had a curve fitted in MATLAB (function F = A(1 -e^-t/T^)). This code (provided as a supplementary file, ‘AJSprocess3b.m’) also generated an amplitude value for the level of expression produced by each reaction. Due to the high level of noise in some plots, secondary code (provided as a supplementary file, ‘fitQC.m’) was also implemented to act as a quality control to determine if the curves and subsequent amplitudes were appropriate to use based on additional criteria. If the fitting was inaccurate, an alternative amplitude was determined by calculating the average and population standard deviation of the last five data points. Each construct had its amplitudes for each pH level normalised to the peak value within that their set, putting every construct on the same scale for easier comparison. Data analysis was performed as described in the text.

## Data and code availability

Data and code are available in the supplementary information attached to this manuscript and/or in Ref. [48].

## Acknowledgements

This research was funded in part by the Wellcome Trust (Grant number 204804/Z/16/Z). We also gratefully acknowledge financial support from the School of Engineering, University of Edinburgh (PhD studentship and new academic start-up fund) and BBSRC (Impact Acceleration Award, University of Edinburgh, reference BBSRC IAA PIII107). We are grateful to the Una Europa alliance for providing funding for AJS to travel to Berlin to present this and other research at a conference in Berlin organised by the unano consortium. We wish to thank the following individuals for useful discussions and/or loan of equipment: Prof Stewart Smith, Dr Jamie Marland, Dr Simone Dimartino, Dr Nacho Tudela-Montes, Dr Nadanai Laohakunakorn, Dr Phil Hands, Prof Chris French, Dr Felipe Milacura and Prof Alistair Elfick. Parts of the research described here were conducted using material produced with the assistance of the Edinburgh Genome Foundry (EGF), an engineering biology research facility specialising in the modular, automated assembly of DNA constructs at the University of Edinburgh. We thank the EGF team for their efforts, particularly Dr Sophie Donovan. For the purpose of open access, the authors have applied a CC BY public copyright licence to any Author Accepted Manuscript version arising from this submission.

## Author contributions

Conceptualization – KED & AJS

Methodology – AJS & KED

Software – primarily AJS, with some assistance from KED

Formal analysis – primarily AJS, with some assistance from KED

Investigation – AJS

Data Curation – AJS

Writing - Original Draft – AJS

Writing - Review & Editing – KED & AJS

Visualization – AJS & KED

Supervision – KED

Project administration – KED

Funding acquisition – KED with assistance from AJS

## Additional information

Supplementary Information is available for this paper.

Correspondence and requests for materials should be addressed to Katherine Dunn.

AI statement: we used the model Claude Sonnet 5 within our university’s AI gateway ELM (https://elm.edina.ac.uk/) to proof-read our manuscript. We gave the model the following prompt: ‘Review this scientific paper and identify (giving the line numbers) any formatting or stylistic issues, particularly including inconsistent capitalisation of terms and punctuation usage. Style conventions should be as used by Springer Nature; British English is used.’ We reviewed the model’s output and made changes to the manuscript accordingly.

