## Supplementary material for "Modulating cell-free transcription electrolytically with switchable DNA triplexes": SI - v4 revised post review.pdf

### 1 Sequences and extinction coefficients

#### 1.1 Sequences

##### 1.1.1 Triplex control testing sequences

Sequences are listed below. Variations that include fluorophore labels are included using the IDT sequence codes for applying modifications. Construct names include their modifications, e.g. Cy5, BHQ2, or UL (Unlabelled). Sequences are shown 5' to 3'.

**Triplex testing sequences (used for experiments presented Fig. 2)**

| **Component** | **Sequence (5' - 3')** |
|---|---|
| TFR10_Cy5 | TTCTTTTCTTGAGCTCACTATGCTTCGTCCTTT GTCTGTCTCTATTCTTTTCTT/3Cy5Sp/ |
| TFR10_UL | TTCTTTTCTTGAGCTCACTATGCTTCGTCCTTT GTCTGTCTCTATTCTTTTCTT |
| C10_Cy5 | TTCTTTTCTTGAGCTCACTATGCTTCGTCCTTT GTCTGTCTCTAGTGTCCGTCC/3Cy5Sp/ |
| C10_UL | TTCTTTTCTTGAGCTCACTATGCTTCGTCCTTT GTCTGTCTCTAGTGTCCGTCC |
| SDR10_BHQ2 | AGTGAGCTCAAGAAAAGAA/3BHQ_2/ |
| SDR10_UL | AGTGAGCTCAAGAAAAGAA |
| SDR_40 | TAATACGACTGAGCTCATAGAGAAAAAAGAAA GAAAAGAA |
| N40 | TTCTTTTCTTTCTTTTTTCTCTATGAGCTCAGT CGTATTATTTTTTTTTTTTTTTTTTTTTTTTTT TTTTTTTTTTTTTTTCTTTTTTCTTTCTTTTCT T |
| C40 | TTCTTTTCTTTCTTTTTTCTCTATGAGCTCAGT CGTATTATCACATCACAGAGGCTTTACTTACT ATCAATCTAACTTTACTACACATCATACACTTC CA |
| T7D_TFR20_N54_ds1_BHQ2 | CCCGCGAAATTAATACGACTCACTATAGGGAA AGAAAAAAAAAAAGAAAA/3BHQ_2/ |
| T7D_TFR20_N54_ds2_Cy5 | AGTCGTATTAATTTCGCGGGGTGGTGGTGTGT CTGGTGTGGTGGTCTGGGTGTGTGGTCTGGGT GTGTGGTCTGTTTCTTTTTTTTTTTCTTTT/3C y5Sp/ |
| T7D_TFR20_N54_ds3 | TTTTCTTTTTTTTTTTCTTTCCCTATAGTG |
| T7D_TFR20_N54_ds1_UL | CCCGCGAAATTAATACGACTCACTATAGGGAA AGAAAAAAAAAAAGAAAA |
| T7P_TFR40_N9_ds1_BHQ2 | CCCGCGAAATTAATACGACTCACTATAGGGAA AGAAAAAAAAAAA/3BHQ_2/ |
| T7P_TFR40_N9_ds2_Cy5 | TTTTTTTTTTTCTTTCCCTATAGTGAGTCGTAT TAATTTCGCGGGGTGTGTTTTCTTTCCTTATG CTTTTTGATATCCCTTTCTTTTTTTTTTT/3Cy5 Sp/ |
| T7P_TFR40_N9_ds1_UL | CCCGCGAAATTAATACGACTCACTATAGGGAA AGAAAAAAAAAAA |

##### 1.1.2 Plasmid sequences

aPCR templates were synthesised as Invitrogen$^{TM}$ GeneArt$^{TM}$ plasmids. They each shared the following main sequences for the T7 promoter, TBR (triplex-binding region), iSpinach, and terminator, with the only differences arising in the TBR between T-AT ratio variants. The sequence of the coding strand in the plasmid is defined by combining T7 promoter - TBR - F30_iSpinach - terminator sequences in that order.

**dsDNA template component sequences**

| **Component** | **Sequence (5' - 3')** |
|---|---|
| T7 Promoter | CCCGCGAAATTAATACGACTCACTATAGGG |
| T7D_90 TBR | AAAGAAAAAAAAAAAGAAAAAAGAAAAAAAAAAAAAAGAA |
| T7D_80 TBR | AAAGAAGAAAAAAAGGAAAAAAGAAAAGAAAAAGAAAGAA |
| T7D_70 TBR | AGAGAAGAAAAAAAGGAAGAAAGAAGAGAAAAGGAAAGAA |
| T7D_60 TBR | AGAGAAGGAAAAAAGGAGGAAAGAAGAGGAAGGGAAAGAA |
| F30_iSpinach | TAGTGCTAGCGAATTCTTGCCATGTGTATGTGGGAGACGCGA<br>CTACGGTGAGGGTCGGGTCCAGTAGCTTCGGCTACTGTTGA<br>GTAGAGTGTGGGCTCCGTAGTCGCGTCTCCCCACATACTCTG<br>ATGATCCTTCGGGATCATTCATGGC |
| T7 Terminator | AAATAACCCCTTGGGGCCTCTAAACGGGTCTTGAGGGGTACT<br>AGT |

The 'top' strands (blue in Fig. 3) formed by PCR/aPCR had the same sequence as the coding strand in the plasmid. The 'bottom strands' had the reverse complement of that sequence, with the addition of an overhanging ssDNA tail (next table) as the active part of the EDGE construct. These tail sequences are shown 5'-3', and are attached to the bottom strand sequences in the following order:
T7 terminator - F30_iSpinach - TBR - T7 Promoter - ssDNA tail.

**T7D_90 triplex-forming ssDNA tail sequences**

| Construct | ssDNA tail sequence (5' - 3') |
|---|---|
| T7D_90_TFR40_N54 | GTGGTGGTGTGTCTGGTGTGGTGGTCTGGGTGTGTGGTCTGGGTGTGTGGTCTGTTTCTTTTTTTTTTTCTTTTTTCTTTTTTTTTTTTTTTCTT |
| T7D_90_TFR40_N48 | GTGGTGGTGTGTCTGGTGTGGTGGTCTGGGTGTGTGGTCTGGGTGTGTTTTCTTTTTTTTTTTTCTTTTTTCTTTTTTTTTTTTTTCTT |
| T7D_90_TFR40_N42 | GTGGTGGTGTGTCTGGTGTGGTGGTCTGGGTGTGTGGTCTGGTTTCTTTTTTTTTTTCTTTTTTCTTTTTTTTTTTTTTTTCTT |
| T7D_90_TFR30_N54 | GTGGTGGTGTGTCTGGTGTGGTGGTCTGGGTGTGTGGTCTGGGTGTGTGGTCTGTTTCTTTTTTTTTTTCTTTTTTCTTTTTTT |
| T7D_90_TFR30_N48 | GTGGTGGTGTGTCTGGTGTGGTGGTCTGGGTGTGTGGTCTGGGTGTGTTTTCTTTTTTTTTTTTCTTTTTTCTTTTTTTT |
| T7D_90_TFR30_N42 | GTGGTGGTGTGTCTGGTGTGGTGGTCTGGGTGTGTGGTCTGGTTTCTTTTTTTTTTTCTTTTTTCTTTTTTTT |
| T7D_90_TFR20_N54 | GTGGTGGTGTGTCTGGTGTGGTGGTCTGGGTGTGTGGTCTGGGTGTGTGGTCTGTTTCTTTTTTTTTTTCTTTT |
| T7D_90_TFR20_N48 | GTGGTGGTGTGTCTGGTGTGGTGGTCTGGGTGTGTGGTCTGGGTGTGTTTTCTTTTTTTTTTTTCTTTT |
| T7D_90_TFR20_N42 | GTGGTGGTGTGTCTGGTGTGGTGGTCTGGGTGTGTGGTCTGGTTTCTTTTTTTTTTTCTTTT |
| T7D_90_TFR10_N54 | GTGGTGGTGTGTCTGGTGTGGTGGTCTGGGTGTGTGGTCTGGGTGTGTGGTCTGTTTCTTTTTT |
| T7D_90_TFR10_N48 | GTGGTGGTGTGTCTGGTGTGGTGGTCTGGGTGTGTGGTCTGGGTGTGTTTTCTTTTTT |
| T7D_90_TFR10_N42 | GTGGTGGTGTGTCTGGTGTGGTGGTCTGGGTGTGTGGTCTGGTTTCTTTTTT |

**T7D_90 control ssDNA tail sequences**

| Construct | ssDNA tail sequence (5' - 3') |
|---|---|
| T7D_90_C40 _N54 | GTGGTGGTGTGTCTGGTGTGGTGGTCTGGGTGTGTGGTCTG GGTGTGTGGTCTGTGGTGGTGGTGTCTGGTGTGTGGTGTGT GGTGGGTGGTCT |
| T7D_90_C40 _N48 | GTGGTGGTGTGTCTGGTGTGGTGGTCTGGGTGTGTGGTCTG GGTGTGTTGGTGGTGGTGTCTGGTGTGTGGTGTGTGGTGGG TGGTCT |
| T7D_90_C40 _N42 | GTGGTGGTGTGTCTGGTGTGGTGGTCTGGGTGTGTGGTCTG GTGGTGGTGGTGTCTGGTGTGTGGTGTGTGGTGGGTGGTCT |
| T7D_90_C30 _N54 | GTGGTGGTGTGTCTGGTGTGGTGGTCTGGGTGTGTGGTCTG GGTGTGTGGTCTGTGGTGGTGGTGTCTGGTGTGTGGTGTGT GG |
| T7D_90_C30 _N48 | GTGGTGGTGTGTCTGGTGTGGTGGTCTGGGTGTGTGGTCTG GGTGTGTTGGTGGTGGTGTCTGGTGTGTGGTGTGTGG |
| T7D_90_C30 _N42 | GTGGTGGTGTGTCTGGTGTGGTGGTCTGGGTGTGTGGTCTG GTGGTGGTGGTGTCTGGTGTGTGGTGTGTGG |
| T7D_90_C20 _N54 | GTGGTGGTGTGTCTGGTGTGGTGGTCTGGGTGTGTGGTCTG GGTGTGTGGTCTGTGGTGGTGGTGTCTGGTGTG |
| T7D_90_C20 _N48 | GTGGTGGTGTGTCTGGTGTGGTGGTCTGGGTGTGTGGTCTG GGTGTGTTGGTGGTGGTGTCTGGTGTG |
| T7D_90_C20 _N42 | GTGGTGGTGTGTCTGGTGTGGTGGTCTGGGTGTGTGGTCTG GTGGTGGTGGTGTCTGGTGTG |
| T7D_90_C10 _N54 | GTGGTGGTGTGTCTGGTGTGGTGGTCTGGGTGTGTGGTCTG GGTGTGTGGTCTGTGGTGGTGGT |
| T7D_90_C10 _N48 | GTGGTGGTGTGTCTGGTGTGGTGGTCTGGGTGTGTGGTCTG GGTGTGTTGGTGGTGGT |
| T7D_90_C10 _N42 | GTGGTGGTGTGTCTGGTGTGGTGGTCTGGGTGTGTGGTCTG GTGGTGGTGGT |

**T7D_80 triplex-forming ssDNA tail sequences**

| **Construct** | **ssDNA tail sequence (5' - 3')** |
|---|---|
| T7D_80_TFR40_N54 | GTGGTGGTGTGTCTGGTGTGGTGGTCTGGGTGTGTGGTCTGGGTGTGTGGTCTGTTTCTTCTTTTTTTCCTTTTTTCTTTTCTTTTTCTTTCTT |
| T7D_80_TFR40_N48 | GTGGTGGTGTGTCTGGTGTGGTGGTCTGGGTGTGTGGTCTGGGTGTGTTTTCTTCTTTTTTTCCTTTTTTCTTTTCTTTTTCTTTCTT |
| T7D_80_TFR40_N42 | GTGGTGGTGTGTCTGGTGTGGTGGTCTGGGTGTGTGGTCTGGTTTCTTCTTTTTTTCCTTTTTTCTTTTCTTTTTCTTTCTT |
| T7D_80_TFR30_N54 | GTGGTGGTGTGTCTGGTGTGGTGGTCTGGGTGTGTGGTCTGGGTGTGTGGTCTGTTTCTTCTTTTTTTCCTTTTTTCTTTTCTT |
| T7D_80_TFR30_N48 | GTGGTGGTGTGTCTGGTGTGGTGGTCTGGGTGTGTGGTCTGGGTGTGTTTTCTTCTTTTTTTCCTTTTTTCTTTTCTT |
| T7D_80_TFR30_N42 | GTGGTGGTGTGTCTGGTGTGGTGGTCTGGGTGTGTGGTCTGGTTTCTTCTTTTTTTCCTTTTTTCTTTTCTT |
| T7D_80_TFR20_N54 | GTGGTGGTGTGTCTGGTGTGGTGGTCTGGGTGTGTGGTCTGGGTGTGTGGTCTGTTTCTTCTTTTTTTCCTTTT |
| T7D_80_TFR20_N48 | GTGGTGGTGTGTCTGGTGTGGTGGTCTGGGTGTGTGGTCTGGGTGTGTTTTCTTCTTTTTTTCCTTTT |
| T7D_80_TFR20_N42 | GTGGTGGTGTGTCTGGTGTGGTGGTCTGGGTGTGTGGTCTGGTTTCTTCTTTTTTTCCTTTT |
| T7D_80_TFR10_N54 | GTGGTGGTGTGTCTGGTGTGGTGGTCTGGGTGTGTGGTCTGGGTGTGTGGTCTGTTTCTTCTTT |
| T7D_80_TFR10_N48 | GTGGTGGTGTGTCTGGTGTGGTGGTCTGGGTGTGTGGTCTGGGTGTGTTTTCTTCTTT |
| T7D_80_TFR10_N42 | GTGGTGGTGTGTCTGGTGTGGTGGTCTGGGTGTGTGGTCTGGTTTCTTCTTT |

**T7D_80 control ssDNA tail sequences**

| Construct | ssDNA tail sequence (5' - 3') |
|---|---|
| T7D_80_C40_N54 | GTGGTGGTGTGTCTGGTGTGGTGGTCTGGGTGTGTGGTCTGGGTGTGTGGTCTGTGGTGGTGGTGTCTGGTGTGTGGTGTGTGGTGGGTGGTCT |
| T7D_80_C40_N48 | GTGGTGGTGTGTCTGGTGTGGTGGTCTGGGTGTGTGGTCTGGGTGTGTTGGTGGTGGTGTCTGGTGTGTGGTGTGTGGTGGGTGGTCT |
| T7D_80_C40_N42 | GTGGTGGTGTGTCTGGTGTGGTGGTCTGGGTGTGTGGTCTGGTGGTGGTGGTGTCTGGTGTGTGGTGTGTGGTGGGTGGTCT |
| T7D_80_C30_N54 | GTGGTGGTGTGTCTGGTGTGGTGGTCTGGGTGTGTGGTCTGGGTGTGTGGTCTGTGGTGGTGGTGTCTGGTGTGTGGTGTGTGG |
| T7D_80_C30_N48 | GTGGTGGTGTGTCTGGTGTGGTGGTCTGGGTGTGTGGTCTGGGTGTGTTGGTGGTGGTGTCTGGTGTGTGGTGTGTGG |
| T7D_80_C30_N42 | GTGGTGGTGTGTCTGGTGTGGTGGTCTGGGTGTGTGGTCTGGTGGTGGTGGTGTCTGGTGTGTGGTGTGTGG |
| T7D_80_C20_N54 | GTGGTGGTGTGTCTGGTGTGGTGGTCTGGGTGTGTGGTCTGGGTGTGTGGTCTGTGGTGGTGGTGTCTGGTGTG |
| T7D_80_C20_N48 | GTGGTGGTGTGTCTGGTGTGGTGGTCTGGGTGTGTGGTCTGGGTGTGTTGGTGGTGGTGTCTGGTGTG |
| T7D_80_C20_N42 | GTGGTGGTGTGTCTGGTGTGGTGGTCTGGGTGTGTGGTCTGGTGGTGGTGGTGTCTGGTGTG |
| T7D_80_C10_N54 | GTGGTGGTGTGTCTGGTGTGGTGGTCTGGGTGTGTGGTCTGGGTGTGTGGTCTGTGGTGGTGGT |
| T7D_80_C10_N48 | GTGGTGGTGTGTCTGGTGTGGTGGTCTGGGTGTGTGGTCTGGGTGTGTTGGTGGTGGT |
| T7D_80_C10_N42 | GTGGTGGTGTGTCTGGTGTGGTGGTCTGGGTGTGTGGTCTGGTGGTGGTGGT |

**T7D_70 triplex-forming ssDNA tail sequences**

| **Construct** | **ssDNA tail sequence (5' - 3')** |
|---|---|
| T7D_70_TFR40_N54 | GTGGTGGTGTGTCTGGTGTGGTGGTCTGGGTGTGTGGTCTGGGTGTGTGGTCTGTCTCTTCTTTTTTTCCTTCTTTCTTCTCTTTTCCTTTCTT |
| T7D_70_TFR40_N48 | GTGGTGGTGTGTCTGGTGTGGTGGTCTGGGTGTGTGGTCTGGGTGTGTTCTCTTCTTTTTTTCCTTCTTTCTTCTCTTTTCCTTTCTT |
| T7D_70_TFR40_N42 | GTGGTGGTGTGTCTGGTGTGGTGGTCTGGGTGTGTGGTCTGGTCTCTTCTTTTTTTCCTTCTTTCTTCTCTTTTCCTTTCTT |
| T7D_70_TFR30_N54 | GTGGTGGTGTGTCTGGTGTGGTGGTCTGGGTGTGTGGTCTGGGTGTGTGGTCTGTCTCTTCTTTTTTTCCTTCTTTCTTCTCTT |
| T7D_70_TFR30_N48 | GTGGTGGTGTGTCTGGTGTGGTGGTCTGGGTGTGTGGTCTGGGTGTGTTCTCTTCTTTTTTTCCTTCTTTCTTCTCTT |
| T7D_70_TFR30_N42 | GTGGTGGTGTGTCTGGTGTGGTGGTCTGGGTGTGTGGTCTGGTCTCTTCTTTTTTTCCTTCTTTCTTCTCTT |
| T7D_70_TFR20_N54 | GTGGTGGTGTGTCTGGTGTGGTGGTCTGGGTGTGTGGTCTGGGTGTGTGGTCTGTCTCTTCTTTTTTTCCTTCT |
| T7D_70_TFR20_N48 | GTGGTGGTGTGTCTGGTGTGGTGGTCTGGGTGTGTGGTCTGGGTGTGTTCTCTTCTTTTTTTCCTTCT |
| T7D_70_TFR20_N42 | GTGGTGGTGTGTCTGGTGTGGTGGTCTGGGTGTGTGGTCTGGTCTCTTCTTTTTTTCCTTCT |
| T7D_70_TFR10_N54 | GTGGTGGTGTGTCTGGTGTGGTGGTCTGGGTGTGTGGTCTGGGTGTGTGGTCTGTCTCTTCTTT |
| T7D_70_TFR10_N48 | GTGGTGGTGTGTCTGGTGTGGTGGTCTGGGTGTGTGGTCTGGGTGTGTTCTCTTCTTT |
| T7D_70_TFR10_N42 | GTGGTGGTGTGTCTGGTGTGGTGGTCTGGGTGTGTGGTCTGGTCTCTTCTTT |

**T7D_70 control ssDNA tail sequences**

| Construct | ssDNA tail sequence (5' - 3') |
|---|---|
| T7D_70_C40 _N54 | GTGGTGGTGTGTCTGGTGTGGTGGTCTGGGTGTGTGGTCTG GGTGTGTGGTCTGTGGTGGTGGTGTCTGGTGTGTGGTGTGT GGTGGGTGGTCT |
| T7D_70_C40 _N48 | GTGGTGGTGTGTCTGGTGTGGTGGTCTGGGTGTGTGGTCTG GGTGTGTTGGTGGTGGTGTCTGGTGTGTGGTGTGTGGTGGG TGGTCT |
| T7D_70_C40 _N42 | GTGGTGGTGTGTCTGGTGTGGTGGTCTGGGTGTGTGGTCTG GTGGTGGTGGTGTCTGGTGTGTGGTGTGTGGTGGGTGGTCT |
| T7D_70_C30 _N54 | GTGGTGGTGTGTCTGGTGTGGTGGTCTGGGTGTGTGGTCTG GGTGTGTGGTCTGTGGTGGTGGTGTCTGGTGTGTGGTGTGT GG |
| T7D_70_C30 _N48 | GTGGTGGTGTGTCTGGTGTGGTGGTCTGGGTGTGTGGTCTG GGTGTGTTGGTGGTGGTGTCTGGTGTGTGGTGTGTGG |
| T7D_70_C30 _N42 | GTGGTGGTGTGTCTGGTGTGGTGGTCTGGGTGTGTGGTCTG GTGGTGGTGGTGTCTGGTGTGTGGTGTGTGG |
| T7D_70_C20 _N54 | GTGGTGGTGTGTCTGGTGTGGTGGTCTGGGTGTGTGGTCTG GGTGTGTGGTCTGTGGTGGTGGTGTCTGGTGTG |
| T7D_70_C20 _N48 | GTGGTGGTGTGTCTGGTGTGGTGGTCTGGGTGTGTGGTCTG GGTGTGTTGGTGGTGGTGTCTGGTGTG |
| T7D_70_C20 _N42 | GTGGTGGTGTGTCTGGTGTGGTGGTCTGGGTGTGTGGTCTG GTGGTGGTGGTGTCTGGTGTG |
| T7D_70_C10 _N54 | GTGGTGGTGTGTCTGGTGTGGTGGTCTGGGTGTGTGGTCTG GGTGTGTGGTCTGTGGTGGTGGT |
| T7D_70_C10 _N48 | GTGGTGGTGTGTCTGGTGTGGTGGTCTGGGTGTGTGGTCTG GGTGTGTTGGTGGTGGT |
| T7D_70_C10 _N42 | GTGGTGGTGTGTCTGGTGTGGTGGTCTGGGTGTGTGGTCTG GTGGTGGTGGT |

**T7D_60 triplex-forming ssDNA tail sequences**

| Construct | ssDNA tail sequence (5' - 3') |
|---|---|
| T7D_60_TFR 40_N54 | GTGGTGGTGTGTCTGGTGTGGTGGTCTGGGTGTGTGGTCTG GGTGTGTGGTCTGTCTCTTCCTTTTTTCCTCCTTTCTTCTCC TTCCCTTTCTT |
| T7D_60_TFR 40_N48 | GTGGTGGTGTGTCTGGTGTGGTGGTCTGGGTGTGTGGTCTG GGTGTGTTCTCTTCCTTTTTTCCTCCTTTCTTCTCCTTCCCTT TCTT |
| T7D_60_TFR 40_N42 | GTGGTGGTGTGTCTGGTGTGGTGGTCTGGGTGTGTGGTCTG GTCTCTTCCTTTTTTCCTCCTTTCTTCTCCTTCCCTTTCTT |
| T7D_60_TFR 30_N54 | GTGGTGGTGTGTCTGGTGTGGTGGTCTGGGTGTGTGGTCTG GGTGTGTGGTCTGTCTCTTCCTTTTTTCCTCCTTTCTTCTCC T |
| T7D_60_TFR 30_N48 | GTGGTGGTGTGTCTGGTGTGGTGGTCTGGGTGTGTGGTCTG GGTGTGTTCTCTTCCTTTTTTCCTCCTTTCTTCTCCT |
| T7D_60_TFR 30_N42 | GTGGTGGTGTGTCTGGTGTGGTGGTCTGGGTGTGTGGTCTG GTCTCTTCCTTTTTTCCTCCTTTCTTCTCCT |
| T7D_60_TFR 20_N54 | GTGGTGGTGTGTCTGGTGTGGTGGTCTGGGTGTGTGGTCTG GGTGTGTGGTCTGTCTCTTCCTTTTTTCCTCCT |
| T7D_60_TFR 20_N48 | GTGGTGGTGTGTCTGGTGTGGTGGTCTGGGTGTGTGGTCTG GGTGTGTTCTCTTCCTTTTTTCCTCCT |
| T7D_60_TFR 20_N42 | GTGGTGGTGTGTCTGGTGTGGTGGTCTGGGTGTGTGGTCTG GTCTCTTCCTTTTTTCCTCCT |
| T7D_60_TFR 10_N54 | GTGGTGGTGTGTCTGGTGTGGTGGTCTGGGTGTGTGGTCTG GGTGTGTGGTCTGTCTCTTCCTT |
| T7D_60_TFR 10_N48 | GTGGTGGTGTGTCTGGTGTGGTGGTCTGGGTGTGTGGTCTG GGTGTGTTCTCTTCCTT |
| T7D_60_TFR 10_N42 | GTGGTGGTGTGTCTGGTGTGGTGGTCTGGGTGTGTGGTCTG GTCTCTTCCTT |

**T7D_60 control ssDNA tail sequences**

| **Construct** | **ssDNA tail sequence (5' - 3')** |
|---|---|
| T7D_60_C40_N54 | GTGGTGGTGTGTCTGGTGTGGTGGTCTGGGTGTGTGGTCTG GGTGTGTGGTCTGTGGTGGTGGTGTCTGGTGTGTGGTGTGT GGTGGGTGGTCT |
| T7D_60_C40_N48 | GTGGTGGTGTGTCTGGTGTGGTGGTCTGGGTGTGTGGTCTG GGTGTGTTGGTGGTGGTGTCTGGTGTGTGGTGTGTGGTGGG TGGTCT |
| T7D_60_C40_N42 | GTGGTGGTGTGTCTGGTGTGGTGGTCTGGGTGTGTGGTCTG GTGGTGGTGGTGTCTGGTGTGTGGTGTGTGGTGGGTGGTCT |
| T7D_60_C30_N54 | GTGGTGGTGTGTCTGGTGTGGTGGTCTGGGTGTGTGGTCTG GGTGTGTGGTCTGTGGTGGTGGTGTCTGGTGTGTGGTGTGT GG |
| T7D_60_C30_N48 | GTGGTGGTGTGTCTGGTGTGGTGGTCTGGGTGTGTGGTCTG GGTGTGTTGGTGGTGGTGTCTGGTGTGTGGTGTGTGG |
| T7D_60_C30_N42 | GTGGTGGTGTGTCTGGTGTGGTGGTCTGGGTGTGTGGTCTG GTGGTGGTGGTGTCTGGTGTGTGGTGTGTGG |
| T7D_60_C20_N54 | GTGGTGGTGTGTCTGGTGTGGTGGTCTGGGTGTGTGGTCTG GGTGTGTGGTCTGTGGTGGTGGTGTCTGGTGTG |
| T7D_60_C20_N48 | GTGGTGGTGTGTCTGGTGTGGTGGTCTGGGTGTGTGGTCTG GGTGTGTTGGTGGTGGTGTCTGGTGTG |
| T7D_60_C20_N42 | GTGGTGGTGTGTCTGGTGTGGTGGTCTGGGTGTGTGGTCTG GTGGTGGTGGTGTCTGGTGTG |
| T7D_60_C10_N54 | GTGGTGGTGTGTCTGGTGTGGTGGTCTGGGTGTGTGGTCTG GGTGTGTGGTCTGTGGTGGTGGT |
| T7D_60_C10_N48 | GTGGTGGTGTGTCTGGTGTGGTGGTCTGGGTGTGTGGTCTG GGTGTGTTGGTGGTGGT |
| T7D_60_C10_N42 | GTGGTGGTGTGTCTGGTGTGGTGGTCTGGGTGTGTGGTCTG GTGGTGGTGGT |

##### 1.1.3 Forward primer sequences

Primer sequences are divided into sections: T-AT Ratio (90%, 80%, 70%, 60%) and triplex-forming / control.

**T7D_90 triplex-forming construct forward primer sequences**

| **Construct** | **Primer sequence (5' - 3')** | **BioPython $T_m$ (°C)** | **NEB$^{TM}$ Q5 $T_m$ (°C)** |
|---|---|---|---|
| T7D_90_TFR40_N54 | AAGAAAAAAAAAAAAAAGAAAAAAGAAAAAAAAAAAGAAACAGAC | 56.5 | 64 |
| T7D_90_TFR40_N48 | AAGAAAAAAAAAAAAAAGAAAAAAGAAAAAAAAAAAGAAAACACA | 56.42 | 64 |
| T7D_90_TFR40_N42 | AAGAAAAAAAAAAAAAAGAAAAAAGAAAAAAAAAAAGAAACCAG | 56.41 | 64 |
| T7D_90_TFR30_N54 | AAAAAAAGAAAAAAGAAAAAAAAAAAGAAACAGACCAC | 56.05 | 64 |
| T7D_90_TFR30_N48 | AAAAAAAGAAAAAAGAAAAAAAAAAAGAAAACACACCC | 56.35 | 64 |
| T7D_90_TFR30_N42 | AAAAAAAGAAAAAAGAAAAAAAAAAAGAAACCAGACC | 55.77 | 64 |
| T7D_90_TFR20_N54 | AAAAGAAAAAAAAAAAGAAACAGACCACACAC | 55.75 | 65 |
| T7D_90_TFR20_N48 | AAAAGAAAAAAAAAAAGAAAACACACCCAGAC | 55.73 | 65 |
| T7D_90_TFR20_N42 | AAAAGAAAAAAAAAAAGAAACCAGACCACACA | 56.41 | 66 |
| T7D_90_TFR10_N54 | AAAAAAGAAACAGACCACACACCCAG | 56.46 | 68 |
| T7D_90_TFR10_N48 | AAAAAAGAAAACACACCCAGACCACA | 56.09 | 67 |
| T7D_90_TFR10_N42 | AAAAAAGAAACCAGACCACACACCC | 56.3 | 68 |

**T7D_control construct forward primer sequences**

| Construct | Primer sequence (5' - 3') | BioPython $T_m$ (°C) | NEB™ Q5 $T_m$ (°C) |
|---|---|---|---|
| T7D_90_C40_N54 | AGACCACCCACCACACACC | 55.84 | 70 |
| T7D_90_C40_N48 | AGACCACCCACCACACACC | 55.84 | 70 |
| T7D_90_C40_N42 | AGACCACCCACCACACACC | 55.84 | 70 |
| T7D_90_C30_N54 | CCACACACCACACACCAGAC | 55.33 | 69 |
| T7D_90_C30_N48 | CCACACACCACACACCAGAC | 55.33 | 69 |
| T7D_90_C30_N42 | CCACACACCACACACCAGAC | 55.33 | 69 |
| T7D_90_C20_N54 | CACACCAGACACCACCACCA | 56.37 | 70 |
| T7D_90_C20_N48 | CACACCAGACACCACCACCA | 56.37 | 70 |
| T7D_90_C20_N42 | CACACCAGACACCACCACCA | 56.37 | 70 |
| T7D_90_C10_N54 | ACCACCACCACAGACCACAC | 56.36 | 70 |
| T7D_90_C10_N48 | ACCACCACCAACACACCCA | 55.64 | 70 |
| T7D_90_C10_N42 | ACCACCACCACCAGACCAC | 55.84 | 70 |

**T7D_80 triplex-forming construct forward primer sequences**

| **Construct** | **Primer sequence (5' - 3')** | **BioPython $T_m$ (°C)** | **NEB™ Q5 $T_m$ (°C)** |
|---|---|---|---|
| T7D_80_TFR40_N54 | AAGAAAGAAAAAGAAAAGAAAAAAGGAAAAAAAGAAGAAA | 56.2 | 64 |
| T7D_80_TFR40_N48 | AAGAAAGAAAAAGAAAAGAAAAAAGGAAAAAAAGAAGAAAA | 56.44 | 64 |
| T7D_80_TFR40_N42 | AAGAAAGAAAAAGAAAAGAAAAAAGGAAAAAAAGAAGAAA | 56.2 | 64 |
| T7D_80_TFR30_N54 | AAGAAAAGAAAAAAGGAAAAAAAGAAGAAACAGAC | 55.61 | 64 |
| T7D_80_TFR30_N48 | AAGAAAAGAAAAAAGGAAAAAAAGAAGAAAACACAC | 56.22 | 65 |
| T7D_80_TFR30_N42 | AAGAAAAGAAAAAAGGAAAAAAAGAAGAAACCAGA | 56.21 | 65 |
| T7D_80_TFR20_N54 | AAAAGGAAAAAAAGAAGAAACAGACCACAC | 55.91 | 66 |
| T7D_80_TFR20_N48 | AAAAGGAAAAAAAGAAGAAAACACACCCAG | 56.08 | 66 |
| T7D_80_TFR20_N42 | AAAAGGAAAAAAAGAAGAAACCAGACCAC | 55.54 | 66 |
| T7D_80_TFR10_N54 | AAAGAAGAAACAGACCACACACCC | 55.44 | 67 |
| T7D_80_TFR10_N48 | AAAGAAGAAAACACACCCAGACCAC | 55.86 | 67 |
| T7D_80_TFR10_N42 | AAAGAAGAAACCAGACCACACACC | 55.44 | 67 |

**T7D_80 control construct forward primer sequences**

| Construct | Primer sequence (5' - 3') | BioPython $T_m$ (°C) | NEB™ Q5 $T_m$ (°C) |
|---|---|---|---|
| T7D_80_C40_N54 | AGACCACCCACCACACACC | 55.84 | 70 |
| T7D_80_C40_N48 | AGACCACCCACCACACACC | 55.84 | 70 |
| T7D_80_C40_N42 | AGACCACCCACCACACACC | 55.84 | 70 |
| T7D_80_C30_N54 | CCACACACCACACACCAGAC | 55.33 | 69 |
| T7D_80_C30_N48 | CCACACACCACACACCAGAC | 55.33 | 69 |
| T7D_80_C30_N42 | CCACACACCACACACCAGAC | 55.33 | 69 |
| T7D_80_C20_N54 | CACACCAGACACCACCACCA | 56.37 | 70 |
| T7D_80_C20_N48 | CACACCAGACACCACCACCA | 56.37 | 70 |
| T7D_80_C20_N42 | CACACCAGACACCACCACCA | 56.37 | 70 |
| T7D_80_C10_N54 | ACCACCACCACAGACCACAC | 56.36 | 70 |
| T7D_80_C10_N48 | ACCACCACCAACACACCCA | 55.64 | 70 |
| T7D_80_C10_N42 | ACCACCACCACCAGACCAC | 55.84 | 70 |

**T7D_70 triplex-forming construct forward primer sequences**

| **Construct** | **Primer sequence (5' - 3')** | **BioPython $T_m$ (°C)** | **NEB™ Q5 $T_m$ (°C)** |
|---|---|---|---|
| T7D_70_TFR40_N54 | AAGAAAGGAAAAGAGAAGAAAGAAGGAAAAAAAG | 55.86 | 65 |
| T7D_70_TFR40_N48 | AAGAAAGGAAAAGAGAAGAAAGAAGGAAAAAAAG | 55.86 | 65 |
| T7D_70_TFR40_N42 | AAGAAAGGAAAAGAGAAGAAAGAAGGAAAAAAAG | 55.86 | 65 |
| T7D_70_TFR30_N54 | AAGAGAAGAAAGAAGGAAAAAAAGAAGAGACAG | 56.35 | 66 |
| T7D_70_TFR30_N48 | AAGAGAAGAAAGAAGGAAAAAAAGAAGAGAACA | 56.06 | 66 |
| T7D_70_TFR30_N42 | AAGAGAAGAAAGAAGGAAAAAAAGAAGAGACC | 56.22 | 66 |
| T7D_70_TFR20_N54 | AGAAGGAAAAAAAGAAGAGACAGACCAC | 55.68 | 66 |
| T7D_70_TFR20_N48 | AGAAGGAAAAAAAGAAGAGAACACACCC | 56.09 | 67 |
| T7D_70_TFR20_N42 | AGAAGGAAAAAAAGAAGAGACCAGACCA | 56.45 | 67 |
| T7D_70_TFR10_N54 | AAAGAAGAGACAGACCACACACCC | 56.5 | 69 |
| T7D_70_TFR10_N48 | AAAGAAGAGAACACACCCAGACCA | 55.85 | 68 |
| T7D_70_TFR10_N42 | AAAGAAGAGACCAGACCACACACC | 56.5 | 69 |

**T7D_70 control construct forward primer sequences**

| Construct | Primer sequence (5' - 3') | BioPython $T_m$ (°C) | NEB$^{TM}$ Q5 $T_m$ (°C) |
|---|---|---|---|
| T7D_70_C40_N54 | AGACCACCCACCACACACC | 55.84 | 70 |
| T7D_70_C40_N48 | AGACCACCCACCACACACC | 55.84 | 70 |
| T7D_70_C40_N42 | AGACCACCCACCACACACC | 55.84 | 70 |
| T7D_70_C30_N54 | CCACACACCACACACCAGAC | 55.33 | 69 |
| T7D_70_C30_N48 | CCACACACCACACACCAGAC | 55.33 | 69 |
| T7D_70_C30_N42 | CCACACACCACACACCAGAC | 55.33 | 69 |
| T7D_70_C20_N54 | CACACCAGACACCACCACCA | 56.37 | 70 |
| T7D_70_C20_N48 | CACACCAGACACCACCACCA | 56.37 | 70 |
| T7D_70_C20_N42 | CACACCAGACACCACCACCA | 56.37 | 70 |
| T7D_70_C10_N54 | ACCACCACCACAGACCACAC | 56.36 | 70 |
| T7D_70_C10_N48 | ACCACCACCAACACACCCA | 55.64 | 70 |
| T7D_70_C10_N42 | ACCACCACCACCAGACCAC | 55.84 | 70 |

**T7D_60 triplex-forming construct forward primer sequences**

| **Construct** | **Primer sequence (5' - 3')** | **BioPython $T_m$ (°C)** | **NEB™ Q5 $T_m$ (°C)** |
|---|---|---|---|
| T7D_60_TFR40_N54 | AAGAAAGGGAAGGAGAAGAAAGGAGG | 56.04 | 68 |
| T7D_60_TFR40_N48 | AAGAAAGGGAAGGAGAAGAAAGGAGG | 56.04 | 68 |
| T7D_60_TFR40_N42 | AAGAAAGGGAAGGAGAAGAAAGGAGG | 56.04 | 68 |
| T7D_60_TFR30_N54 | AGGAGAAGAAAGGAGGAAAAAAGGAAGA | 56 | 67 |
| T7D_60_TFR30_N48 | AGGAGAAGAAAGGAGGAAAAAAGGAAGA | 56 | 67 |
| T7D_60_TFR30_N42 | AGGAGAAGAAAGGAGGAAAAAAGGAAGA | 56 | 67 |
| T7D_60_TFR20_N54 | AGGAGGAAAAAAGGAAGAGACAGACC | 56.3 | 68 |
| T7D_60_TFR20_N48 | AGGAGGAAAAAAGGAAGAGAACACAC | 55.33 | 67 |
| T7D_60_TFR20_N42 | AGGAGGAAAAAAGGAAGAGACCAGAC | 56.3 | 68 |
| T7D_60_TFR10_N54 | AAGGAAGAGACAGACCACACACC | 56.09 | 69 |
| T7D_60_TFR10_N48 | AAGGAAGAGAACACACCCAGACC | 56.07 | 69 |
| T7D_60_TFR10_N42 | AAGGAAGAGACCAGACCACACAC | 56.09 | 69 |

**T7D_60 control construct forward primer sequences**

| **Construct** | **Primer sequence (5' - 3')** | **BioPython $T_m$ (°C)** | **NEB™ Q5 $T_m$ (°C)** |
|---|---|---|---|
| T7D_60_C40_N54 | AGACCACCCACCACACACC | 55.84 | 70 |
| T7D_60_C40_N48 | AGACCACCCACCACACACC | 55.84 | 70 |
| T7D_60_C40_N42 | AGACCACCCACCACACACC | 55.84 | 70 |
| T7D_60_C30_N54 | CCACACACCACACACCAGAC | 55.33 | 69 |
| T7D_60_C30_N48 | CCACACACCACACACCAGAC | 55.33 | 69 |
| T7D_60_C30_N42 | CCACACACCACACACCAGAC | 55.33 | 69 |
| T7D_60_C20_N54 | CACACCAGACACCACCACCA | 56.37 | 70 |
| T7D_60_C20_N48 | CACACCAGACACCACCACCA | 56.37 | 70 |
| T7D_60_C20_N42 | CACACCAGACACCACCACCA | 56.37 | 70 |
| T7D_60_C10_N54 | ACCACCACCACAGACCACAC | 56.36 | 70 |
| T7D_60_C10_N48 | ACCACCACCAACACACCCA | 55.64 | 70 |
| T7D_60_C10_N42 | ACCACCACCACCAGACCAC | 55.84 | 70 |

##### 1.1.4 aPCR product custom extinction coefficients

To improve the accuracy of UV-vis quantification of ssDNA, the structures of each aPCR product were simulated using NUPACK to calculate the percentage of ssDNA vs dsDNA. This was then used to calculate a custom extinction coefficient based on extinction coefficients of 0.027$(ng/\mu L)^{-1}cm^{-1}$ for ssDNA and 0.02$(ng/\mu L)^{-1}$ $cm^{-1}$ for dsDNA.

**Top strand aPCR products**

| **Construct** | **Bases paired at 25°C (Ratio)** | **Estimated Extinction Coefficient ($(ng/\mu L)^{-1}cm^{-1}$)** |
|---|---|---|
| Top_Strand_90 | 0.5454668 | 0.023181732 |
| Top_Strand_80 | 0.5473097 | 0.023168832 |
| Top_Strand_70 | 0.5575528 | 0.02309713 |
| Top_Strand_60 | 0.5603459 | 0.023077579 |

**T7D_90 aPCR products**

| **Construct** | **Bases paired at 25°C (Ratio)** | **Estimated Extinction Coefficient ((ng/µL)$^{-1}$cm$^{-1}$)** |
|---|---|---|
| T7D_90_TFR40_N54 | 0.4015855 | 0.024188902 |
| T7D_90_TFR40_N48 | 0.4038156 | 0.024173291 |
| T7D_90_TFR40_N42 | 0.407511 | 0.024147423 |
| T7D_90_TFR30_N54 | 0.4131206 | 0.024108156 |
| T7D_90_TFR30_N48 | 0.4155999 | 0.024090801 |
| T7D_90_TFR30_N42 | 0.4196181 | 0.024062673 |
| T7D_90_TFR20_N54 | 0.4253275 | 0.024022708 |
| T7D_90_TFR20_N48 | 0.4280687 | 0.024003519 |
| T7D_90_TFR20_N42 | 0.4324376 | 0.023972937 |
| T7D_90_TFR10_N54 | 0.4382716 | 0.023932099 |
| T7D_90_TFR10_N48 | 0.4412626 | 0.023911162 |
| T7D_90_TFR10_N42 | 0.4460206 | 0.023877856 |
| T7D_90_C40_N54 | 0.4658456 | 0.023739081 |
| T7D_90_C40_N48 | 0.4739657 | 0.02368224 |
| T7D_90_C40_N42 | 0.460375 | 0.023777375 |
| T7D_90_C30_N54 | 0.4669998 | 0.023731001 |
| T7D_90_C30_N48 | 0.4634333 | 0.023755967 |
| T7D_90_C30_N42 | 0.467535 | 0.023727255 |
| T7D_90_C20_N54 | 0.4465638 | 0.023874053 |
| T7D_90_C20_N48 | 0.4396986 | 0.02392211 |
| T7D_90_C20_N42 | 0.4451754 | 0.023883772 |
| T7D_90_C10_N54 | 0.4487724 | 0.023858593 |
| T7D_90_C10_N48 | 0.444972 | 0.023885196 |
| T7D_90_C10_N42 | 0.4554014 | 0.02381219 |

**T7D_80 aPCR products**

| **Construct** | **Bases paired at 25°C (Ratio)** | **Estimated Extinction Coefficient ((ng/μL)$^{-1}$cm$^{-1}$)** |
|---|---|---|
| T7D_80_TFR40_N54 | 0.4009594 | 0.024193284 |
| T7D_80_TFR40_N48 | 0.4044352 | 0.024168954 |
| T7D_80_TFR40_N42 | 0.4090466 | 0.024136674 |
| T7D_80_TFR30_N54 | 0.4124784 | 0.024112651 |
| T7D_80_TFR30_N48 | 0.4162438 | 0.024086293 |
| T7D_80_TFR30_N42 | 0.4212159 | 0.024051489 |
| T7D_80_TFR20_N54 | 0.4246189 | 0.024027668 |
| T7D_80_TFR20_N48 | 0.4286657 | 0.02399934 |
| T7D_80_TFR20_N42 | 0.4340517 | 0.023961638 |
| T7D_80_TFR10_N54 | 0.4365829 | 0.02394392 |
| T7D_80_TFR10_N48 | 0.4401608 | 0.023918874 |
| T7D_80_TFR10_N42 | 0.4446509 | 0.023887444 |
| T7D_80_C40_N54 | 0.465202 | 0.023743586 |
| T7D_80_C40_N48 | 0.4734132 | 0.023686108 |
| T7D_80_C40_N42 | 0.4816551 | 0.023628414 |
| T7D_80_C30_N54 | 0.4662357 | 0.02373635 |
| T7D_80_C30_N48 | 0.4625965 | 0.023761825 |
| T7D_80_C30_N42 | 0.4660547 | 0.023737617 |
| T7D_80_C20_N54 | 0.4451569 | 0.023883902 |
| T7D_80_C20_N48 | 0.4385551 | 0.023930114 |
| T7D_80_C20_N42 | 0.4441492 | 0.023890956 |
| T7D_80_C10_N54 | 0.4471852 | 0.023869704 |
| T7D_80_C10_N48 | 0.4437137 | 0.023894004 |
| T7D_80_C10_N42 | 0.4541261 | 0.023821117 |

**T7D_70 aPCR products**

| Construct | Bases paired at 25°C (Ratio) | Estimated Extinction Coefficient $((ng/\mu L)^{-1}cm^{-1})$ |
|---|---|---|
| T7D_70_TFR40_N54 | 0.396971 | 0.024221203 |
| T7D_70_TFR40_N48 | 0.3989874 | 0.024207088 |
| T7D_70_TFR40_N42 | 0.4029226 | 0.024179542 |
| T7D_70_TFR30_N54 | 0.4081653 | 0.024142843 |
| T7D_70_TFR30_N48 | 0.4100936 | 0.024129345 |
| T7D_70_TFR30_N42 | 0.4143667 | 0.024099433 |
| T7D_70_TFR20_N54 | 0.4199173 | 0.024060579 |
| T7D_70_TFR20_N48 | 0.4221231 | 0.024045138 |
| T7D_70_TFR20_N42 | 0.4267643 | 0.02401265 |
| T7D_70_TFR10_N54 | 0.4321153 | 0.023975193 |
| T7D_70_TFR10_N48 | 0.4335806 | 0.023964936 |
| T7D_70_TFR10_N42 | 0.4373662 | 0.023938437 |
| T7D_70_C40_N54 | 0.4664144 | 0.023735099 |
| T7D_70_C40_N48 | 0.4753462 | 0.023672577 |
| T7D_70_C40_N42 | 0.4825127 | 0.023622411 |
| T7D_70_C30_N54 | 0.4678122 | 0.023725315 |
| T7D_70_C30_N48 | 0.4651938 | 0.023743643 |
| T7D_70_C30_N42 | 0.4669681 | 0.023731223 |
| T7D_70_C20_N54 | 0.4449734 | 0.023885186 |
| T7D_70_C20_N48 | 0.439718 | 0.023921974 |
| T7D_70_C20_N42 | 0.4440126 | 0.023891912 |
| T7D_70_C10_N54 | 0.4458744 | 0.023878879 |
| T7D_70_C10_N48 | 0.4422063 | 0.023904556 |
| T7D_70_C10_N42 | 0.4525684 | 0.023832021 |

**T7D_60 aPCR products**

| Construct | Bases paired at 25°C (Ratio) | Estimated Extinction Coefficient $((ng/\mu L)^{-1}cm^{-1})$ |
|---|---|---|
| T7D_60_TFR40_N54 | 0.4615594 | 0.023769084 |
| T7D_60_TFR40_N48 | 0.4604648 | 0.023776746 |
| T7D_60_TFR40_N42 | 0.4603041 | 0.023777871 |
| T7D_60_TFR30_N54 | 0.4762404 | 0.023666317 |
| T7D_60_TFR30_N48 | 0.4764878 | 0.023664585 |
| T7D_60_TFR30_N42 | 0.4788909 | 0.023647764 |
| T7D_60_TFR20_N54 | 0.4905981 | 0.023565813 |
| T7D_60_TFR20_N48 | 0.4915699 | 0.023559011 |
| T7D_60_TFR20_N42 | 0.4946373 | 0.023537539 |
| T7D_60_TFR10_N54 | 0.5045351 | 0.023468254 |
| T7D_60_TFR10_N48 | 0.5074946 | 0.023447538 |
| T7D_60_TFR10_N42 | 0.5127601 | 0.023410679 |
| T7D_60_C40_N54 | 0.4730846 | 0.023688408 |
| T7D_60_C40_N48 | 0.4745942 | 0.023677841 |
| T7D_60_C40_N42 | 0.4908136 | 0.023564305 |
| T7D_60_C30_N54 | 0.479689 | 0.023642177 |
| T7D_60_C30_N48 | 0.4823034 | 0.023623876 |
| T7D_60_C30_N42 | 0.5012829 | 0.02349102 |
| T7D_60_C20_N54 | 0.4926643 | 0.02355135 |
| T7D_60_C20_N48 | 0.495488 | 0.023531584 |
| T7D_60_C20_N42 | 0.5151517 | 0.023393938 |
| T7D_60_C10_N54 | 0.5022786 | 0.02348405 |
| T7D_60_C10_N48 | 0.5070003 | 0.023450998 |
| T7D_60_C10_N42 | 0.5280961 | 0.023303327 |

### 2 Circuitry

#### 2.1 Design of circuit for simultaneous electrolytic decomposition in all wells, with constant current

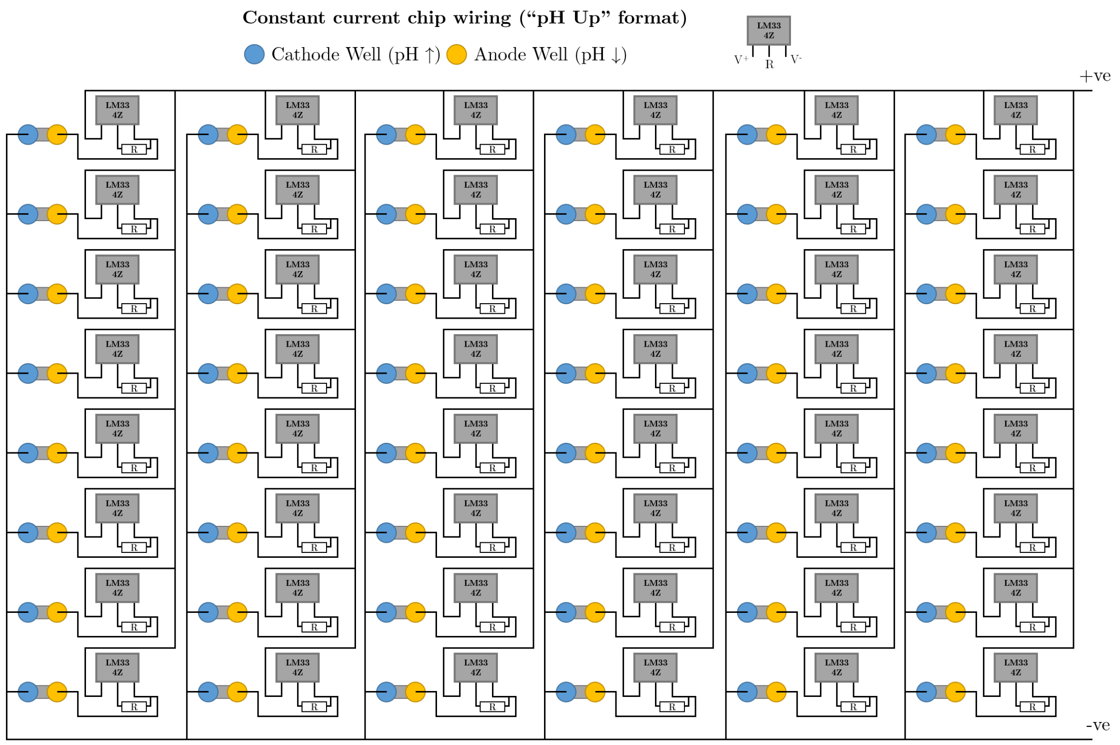


Figure 1: LM334Z constant current chip wiring diagram for connecting EDGE plate wells and electrodes. Electrodes are wired in parallel but include a constant current chip and resistor before each positive electrode. These chips use the resistor to calibrate a set current output, allowing the wells to all be connected in parallel to a single voltage source but to each have the same current flow for electrolysing the EDGE DNA and AceOTB solution. This layout raises the pH in the left hand well. To lower the pH the chip and resistor must be placed on the other side, connected to the left hand electrode and the polarity reversed.

#### 2.2 Photographs of constant current circuitry

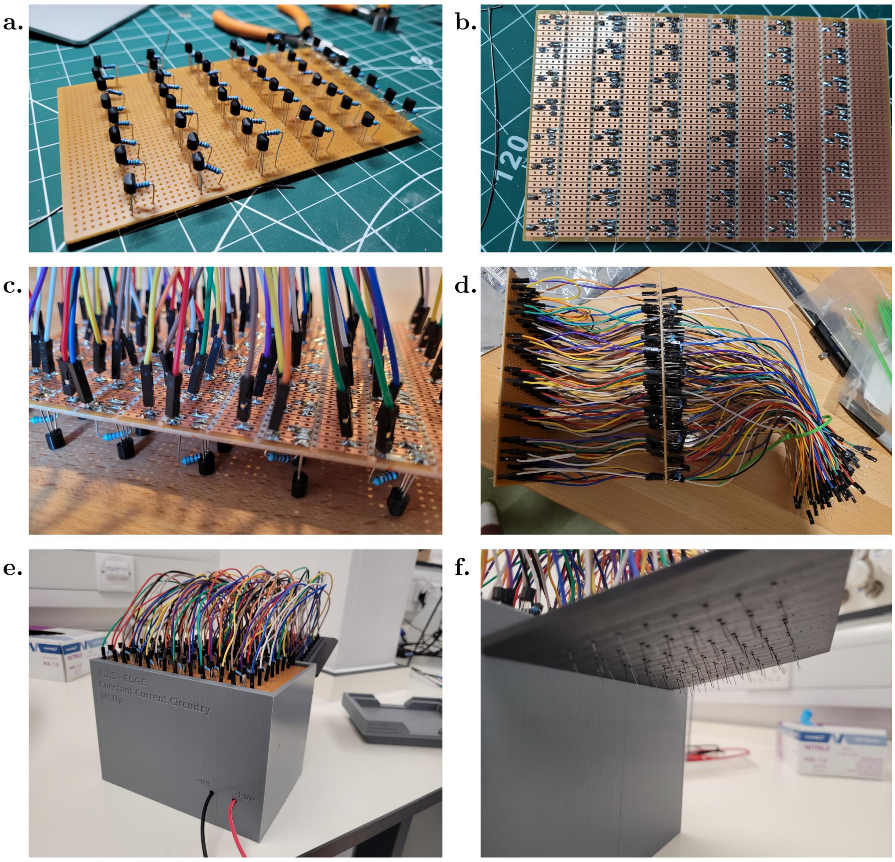

Figure 2: Assembly of whole plate electrolysis circuitry. a) Installation of resistors and LM334Z chips in a regular repeating pattern across stripboard. Each resistor / chip pair corresponds to an individual plate well pair. b) Underside of stripboard. The stripboard is divided into columns, with 12 columns, half for installing resistors and chips and connecting to the positive electrodes, and the other half for connecting the negative electrodes. c) Soldering of Dupont connectors, linking the top stripboard to the bottom stripboard to connect each individual circuit in parallel. d) Completed circuitry, now including longer Dupont connectors that extend away from the stripboard and include platinum electrodes to be glued into the electrode top plate. e) Completely assembled circuitry box (pH up variant). f) Platinum electrodes sticking out of the bottom of the completed electrode top plate.

### 3 Data and code

Alongside this document we provide various supplementary data and code files, as described below.

#### 3.1 Data: EDGE Kinetics Master Sheet

This is an Excel file with multiple tabs. Data comprises baseline corrected averages and population standard deviations from 3 replicates over time, with some of the outliers removed by the original compiler python script used to assemble data from across 3 separate plates. Electrical data is baseline-corrected to 40 minutes, as electrolysis was performed at 40 minutes.

#### 3.2 Code: MATLAB

##### 3.2.1 Code: MATLAB$^{\mathrm{TM}}$ kinetics fitting code

The MATLAB$^{\mathrm{TM}}$ routine provided in the file AJSprocess3b.m is a representative example of code used for curve fitting for iSpinach transcription kinetics results. Prior to use of this code the kinetics data was processed as follows: the baseline was corrected, the average and population standard deviation were calculated, and outliers were removed. Data was processed for 24 constructs at a time (each plate contained 48 constructs, 24 triplex and 24 control, this fitted each set separately for each experiment). The function used was $F = A(1 - e^{-\mathrm{t/T}})$ and fitting was performed using a least-squares algorithm. The parameters $A$ and $T$ were extracted, with the associated standard errors, and exported as a separate excel file.

##### 3.2.2 MATLAB$^{\mathrm{TM}}$ kinetics fitting quality control code

This code (provided in the file FitQC.m) was used to check if the fitting produced by the MATLAB$^{\mathrm{TM}}$ code was sensible, as many were not due to the noise of the results. This code compared the fit to original data for 24 constructs at a time, and used the following criteria to judge each:

1. Is the fractional error on each fit parameter smaller than 50%? If this test is failed it indicates that the error on the fit parameters is excessive.

2. Is the maximum value in the dataset larger than 150? If this test is failed it indicates that there was very little expression.

3. Is the time constant longer than 4 minutes (the minimum spacing between points) and shorter than 360 minutes (three times the duration of the experiment)? If this test is failed it indicates that the kinetics are rising too quickly/slowly to be captured adequately in this experiment.

To be successful each fitting must answer yes to each question. If the fit and/or data do not pass the QC measures, an alternative measure of expression is computed by averaging the last five data points. The error is taken to be the population standard deviation of the five data points in question.

#### 3.3 Code: Python

##### 3.3.1 Primer design

We used a customised python script to design primers, and this is provided as PrimerDesigner1.py.

##### 3.3.2 Compiling kinetic data

To combine the data from the different Overload files produced by our laboratory apparatus, we used a customised python script. This is provided as 'EDWIN Triple Plate Compiler Baseline Corrected'. As the filename suggests, this script also corrected for baseline and removed some outlier data points.
