## Supplementary material for "Modulating cell-free transcription electrolytically with switchable DNA triplexes": Supplementary Note.docx

**Supplementary Note: Incompatibility of DFHBI-based fluorescent RNA aptamers with particular commercial cell-free expression systems**

Alexander J. Speakman & Katherine E. Dunn

School of Engineering, Institute for Bioengineering, University of Edinburgh, The King’s Buildings, Edinburgh, EH9 3DW, Scotland, UK

*This supplementary note is a slightly modified form of the following BioRxiv preprint:* <https://www.biorxiv.org/content/10.1101/2021.08.10.455838v1>

INTRODUCTION:

Fluorophore-binding RNA aptamers are effective tools for investigating RNA synthesis and function.^1^ They consist of single stranded RNA sequences that have been selectively evolved and designed to bind and activate a fluorophore in their solution^2^, which can then be detected by fluorescence spectrophotometry or fluorescence microscopy.^3^ These sequences can be used as a direct fluorescent reporter of transcription without the need for translation^4^, or can be included as a tag on other RNA transcripts and visualised *in vivo* to locate an RNA molecule within a live cell.^5, 6^

Commonly used examples of fluorophore-binding RNAs include Broccoli^7^ and Spinach,^8^ which use 3,5-difluoro-4-hydroxybenzylidene imidazolinone (DFHBI) as a fluorophore.^9^ Once transcribed, Broccoli and Spinach fold to form G-quadruplex structures that allow them to interact with DFHBI.^10^ DFHBI is non-toxic and has a structure that mimics that of the fluorophore within Green Fluorescent Protein (GFP).^11^ As such, DFHBI is non-fluorescent when unbound, but upon binding to RNA aptamers such as Broccoli or Spinach, has a fluorescent activity similar to that of GFP.^12^ There are many other varieties of fluorophore-binding RNA aptamers such as Mango, which binds TO1-Biotin^13^, Pepper, which binds HBC^14^, and Corn, which binds DFHO.^15^ Having these various options makes fluorophore-binding RNA aptamers a versatile tool, offering different wavelengths for excitation and emission, different RNA lengths and structures, and different properties such as photostability and sensitivities to ion concentrations.^16^

As fluorescence directly correlates with the quantity of RNA-fluorophore complexes, RNA aptamers can be used to quantify gene expression, both *in vivo* and *in vitro* when used in cell-free expression systems.^3^ Cell free expression systems are solutions made from purified or reconstituted cell extracts and biological components (E.g, RNA polymerases, ribosomes, etc.).^17, 18^ These allow for a gene to be expressed *in vitro*, by transcribing and translating a desired gene from provided DNA.^19^ However, we have discovered that some commercial cell free expression systems are incompatible for use with some fluorophore-binding RNA aptamers. We attempted to synthesise and detect two different DFHBI-binding RNA aptamers using Invitrogen’s Expressway™ cell-free expression system, which is based on *E. coli* extract and T7 polymerase. These aptamers were iSpinach, a more stable and brighter derivative of Spinach optimised for *in vitro* synthesis^20^, and Broccoli. We found that we were unable to detect any fluorescence produced by our RNA aptamers when using Expressway™. We confirmed that RNA was being produced, and demonstrated that both aptamers do display fluorescent activity when made in a different buffer, but we were unable to make either iSpinach or Broccoli that functioned as intended when using Expressway™.

METHODS:

Linear DNA templates were designed to contain a T7 promoter, a downstream fluorophore-binding RNA aptamer sequence, and a T7 terminator region (see later section for sequences), which were supplied by Integrated DNA Technologies (IDT). As a non-fluorescent control, 500ng of T7 calmodulin plasmid (Invitrogen, #454002) was also used in the following cell free expression reactions. Linear DNA templates from IDT were amplified by PCR using Q5 High fidelity DNA polymerase (NEB, #M0491S) and dNTPs (NEB, #N0447S), before being purified using a Qiaquick PCR purification kit (Qiagen, #28104). PCR was performed using primers from IDT (see later section for sequences), starting with approximately 100ng of DNA amplified for 16 cycles. With the exception of the calmodulin control, approximately 2000ng of DNA template was included in each reaction.

100 µL Expressway™ mini cell-free synthesis kit (Invitrogen, #K990100) reactions were set up in accordance with the user manuals and literature provided by Invitrogen and supplemented with DFHBI (Sigma, #SML1627-5MG) which was dissolved in DMSO (Fisher, #D/4121PB08) and included in the Expressway™ reaction solution at a concentration of 40µM. A previously optimised RNA aptamer transcription buffer (OTB)^21^ modified to be acetate based (AceOTB) was also tested. AceOTB was produced as a 10X stock containing 63mM MgCl­_2_•6(H_2_O) (Sigma, #M2670), 100mM DTT (Melford, #MB1015), 400mM Tris Base (Sigma, #T1503), 20mM Spermidine (Alfa Aesar, #A19096.03) and adjusted to pH 7.5 using 3M Acetic Acid (ACROS Organics, #423225000). This was then diluted to 1X and supplemented with DNA template, rNTPs (2.5mM of each) (NEB, #N0466S), 200U (4µL) of T7 RNA polymerase (NEB, #M0251S), and 40µM DFHBI to the final volume of 100µL. Additional reactions were performed with both Expressway™ and AceOTB that included 100U (5µL) of SUPERase RNase inhibitor (Invitrogen, #AM2696) to prevent any potential RNA degradation that could interfere with aptamer fluorescence. Reactions were performed at room temperature, and upon assembly were loaded into microcuvettes (BioRad, #1702416) positioned in a custom, 3D printed cuvette size adapter (design and photos in supplementary information), and placed in an Agilent Cary Eclipse fluorescence spectrophotometer to detect fluorescence over time (excitation wavelength = 480nm, emission wavelength = 520nm, scanning every 30 seconds). Reactions were scanned for 24 hours to allow for RNA aptamer synthesis.

RNA purification was performed using a Reliaprep™ RNA Miniprep System (Promega, #Z6010), according to the included user manual, with Expressway™ reactions as a source of RNA. For use with the Reliaprep™ kit, individual Expressway™ reactions were set up using 150ng and 1200ng of T7-Broccoli template DNA, and 500ng of calmodulin control plasmid. These were prepared in accordance with the Expressway™ user manuals to a total volume of 50µL and incubated at 37°C for 3 hours, before using the entire volume for purification. Purified RNA was then measured and quantified using a Nanodrop™ UV-visible spectrophotometer (Thermofisher, #ND-2000C) on pedestal mode using 2µL of purified product. Each Nanodrop™ measurement was repeated in triplicate.

RESULTS AND DISCUSSION:

Using Expressway™ to synthesise either DFHBI based fluorophore-binding RNA aptamer yielded no apparent fluorescence. Both iSpinach (*Figure 1a*) and Broccoli (*Figure 1b*) produce fluorescence in the dedicated RNA aptamer AceOTB, but neither aptamer appears to produce fluorescence in Expressway™. A low level of background fluorescence is observed in the Expressway buffer when DFHBI is present, yet it is substantially lower than the fluorescence produced by the same aptamers in AceOTB. It also shows no increase over time as would be expected with cumulative RNA aptamer synthesis. Based on these results it seems that Expressway™ is not suitable for DFHBI based aptamer synthesis. The inclusion of SUPERase also has no effect on the Expressway™ reactions, while its effect in AceOTB is negligible. SUPERase was included to test whether RNAse contamination or endogenous RNases within the *E.coli* extract were degrading the RNA aptamers, however this does not appear to be the issue.

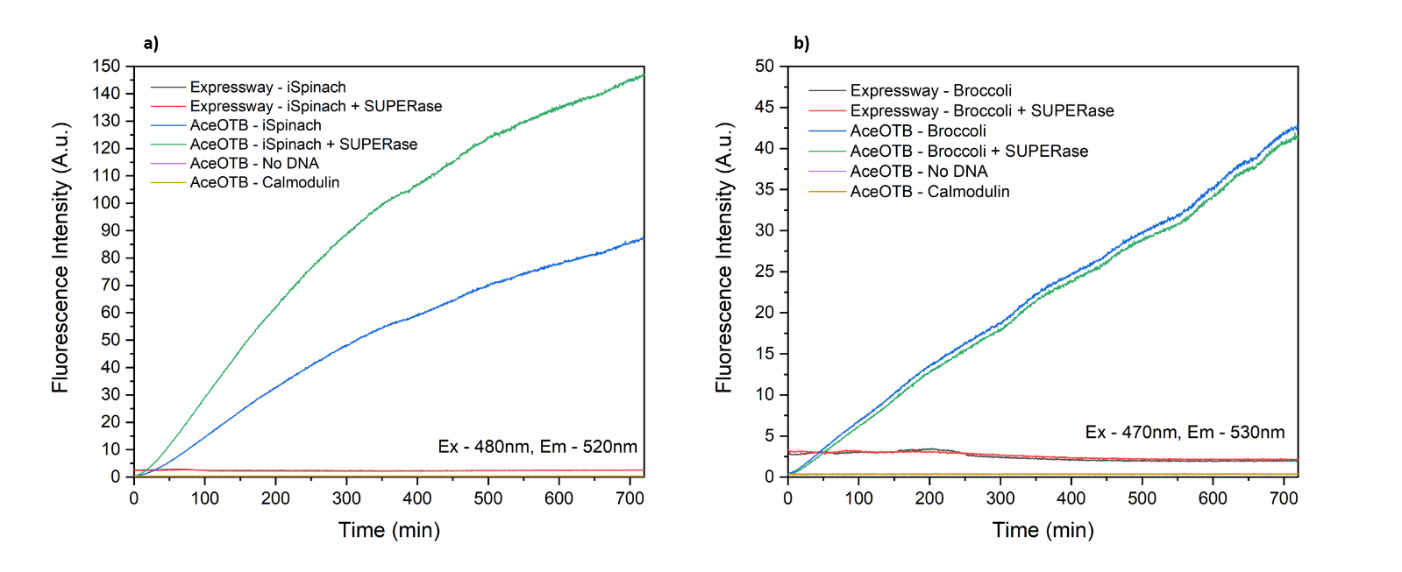

***Figure 1: Fluorescence produced by synthesis of DFHBI-binding RNA aptamer, with 2000ng of DNA template.*** *Note different y-axis scales. Both graphs show fluorescence over time in Expressway™ solution and in AceOTB in vitro transcription buffer, both with and without RNase inhibitor (SUPERase). Both graphs also feature control runs with no DNA template added and a non-fluorescent RNA template (Calmodulin). (a) Fluorescence produced by iSpinach transcription.(b) Fluorescence produced by Broccoli transcription.*

To investigate if the lack of detectable fluorescence was due to insufficient RNA being synthesised, Expressway™ reactions were set up and purified to extract any RNA produced (*Figure 2b)*. RNA purification results showed that a low starting mass of Broccoli aptamer template DNA (150ng) yielded negligible amounts of RNA. Including a larger quantity of DNA (1200ng) produced greater quantities of RNA (*Figure 2b*) even more than the calmodulin control plasmid. The RNA purification data demonstrates that the Expressway mixture is able to transcribe RNA from the Calmodulin control, and that while starting DNA quantity heavily affects RNA yield, the amount of DNA aptamer template required to produce detectable levels on par with the control were well below those used included in the fluorescence spectrophotometry experiments (2000ng of template DNA). Consequently, the issue is not due to an inability to produce Broccoli or iSpinach RNA.

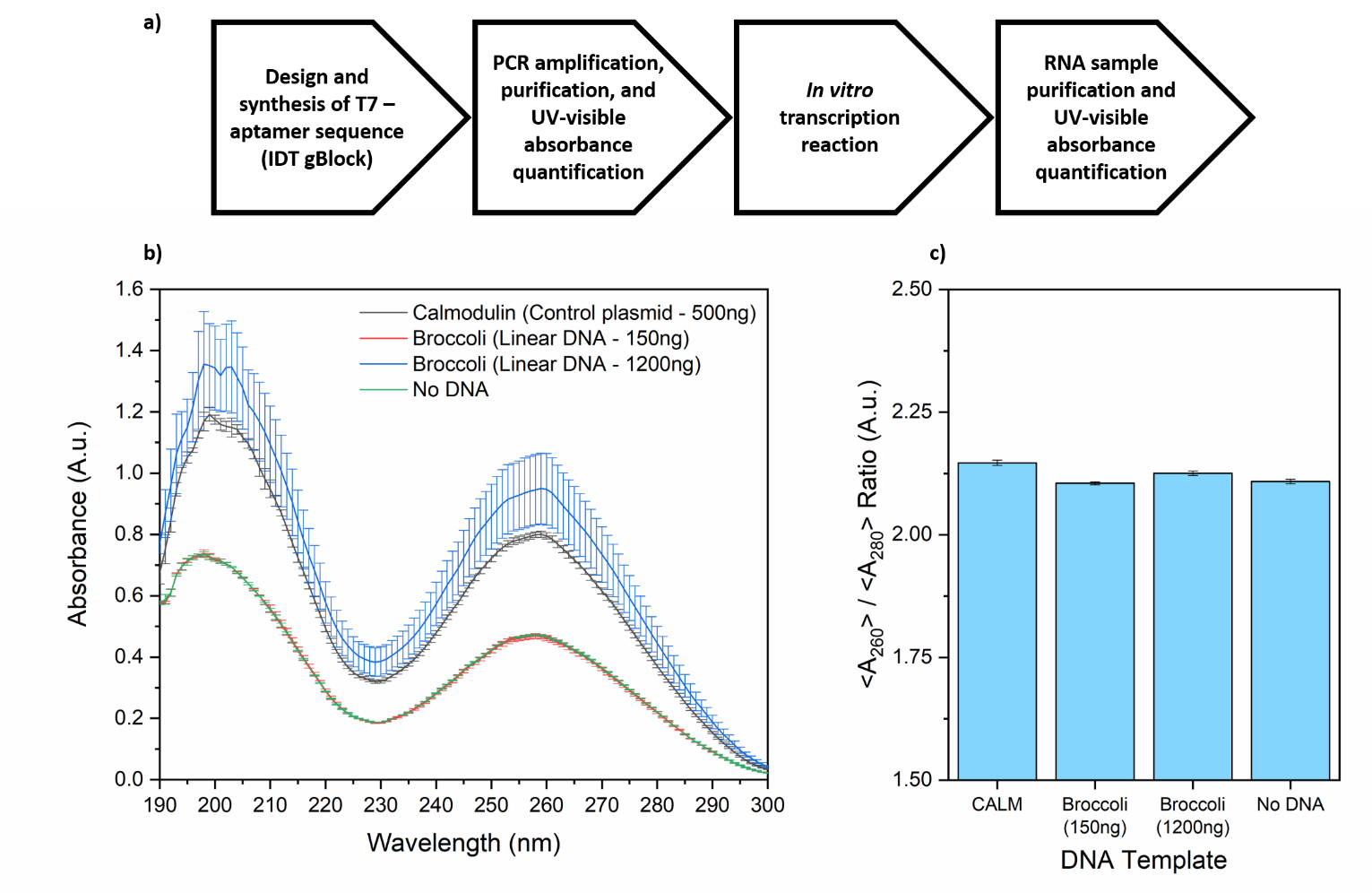

***Figure 2: UV-visible spectroscopy of RNA purification products****. UV-visible absorbance was used to quantify the amount and relative purity of RNA produced by Expressway™ reactions using different templates after purification. (a) Workflow of RNA production from DNA template. (b) Absorbance spectra of purified RNA from each template. (c) Ratio of absorbance at 260nm / 280nm for RNA purified from each template. Absorbance measurements were repeated in triplicate.*

The reason for this lack of fluorescence is unclear, but based on the low-level fluorescence that is present in the DFHBI supplemented Expressway™, and not in the optimised RNA aptamer buffer, one could speculate that a component of Expressway™ is binding the DFHBI in a way that prevents RNA aptamers from binding to produce optimal fluorescence. Given that Expressway™ is primarily used for protein synthesis, it could also be that additional components required during protein synthesis could be binding to the RNA aptamers in a way that prevents the folding and secondary RNA structures required to bind DFHBI. However, as DFHBI-binding aptamers are functional *in vivo* in E.coli^7^, it is difficult to say with certainty which mechanism may be preventing their fluorescence in Expressway™.

This issue does not appear to be limited to Expressway™ cell-free expression systems. It has been reported that a similar lack of fluorescence occurs when attempting to synthesise Spinach2 aptamers using the Promega™ *E.*coli S30™ extract system, using a prokaryotic polymerase and DFHBI-1T, a brighter derivative of DFHBI.^22^ This suggests that the problem is not limited to systems using any particular polymerase, Spinach aptamer variant, or fluorophore derivative, but is potentially caused by something more fundamental in some commercial cell-free expression solutions.

**DNA Sequences (5’ – 3’)**

T7_F30_Broccoli_Term

GTTAGAGAATAGGGATCCGGCGATGCCCGCGAAATTAATACGACTCACTATAGGGTTGCCATGTGTATGTGGGAGACGGTCGGGTCCAGATATTCGTATCTGTCGAGTAGAGTGTGGGCTCCCACATACTCTGATGATCCTTCGGGATCATTCATGGCAAGCACCGCTGAGCAATAACTAGCATAACCCCTTGGGGCCTCTAAACGGGTCTTGAGGGGTTTTTTGAATCTGCAGTATGAATTTACTTTAGGCAACTAGAAGGCACAGT

FWD_Primer_T7_F30_Broccoli

AGAATAGGGATCCGGCGAT

REV_Primer_T7_F30_Broccoli

GCCTAAAGTAAATTCATACTGCAGATTC

T7_F30_iSpinach_Term

GTTAGAGAATAGGGATCCGGAATTCGCGGCCGCTTCTAGAGTAATACGACTCACTATAGGGTTGCCATGTGTATGTGGGAGACGCGACTACGGTGAGGGTCGGGTCCAGTAGCTTCGGCTACTGTTGAGTAGAGTGTGGGCTCCGTAGTCGCGTCTCCCCACATACTCTGATGATCCTTCGGGATCATTCATGGCAAATAACCCCTTGGGGCCTCTAAACGGGTCTTGAGGGGTACTAGTAGCGGCCGCTGCAGGAATCTGCAGTATGAATTTACTTTAGGCAACTAGAAGGCACAGT

FWD_Primer_T7_F30_iSpinach

GCCGCTTCTAGAGTAATACGAC

REV_Primer_T7_F30_iSpinach

GCCTTCTAGTTGCCTAAAGTAAATTC

**3D Printed Microcuvette adapters**
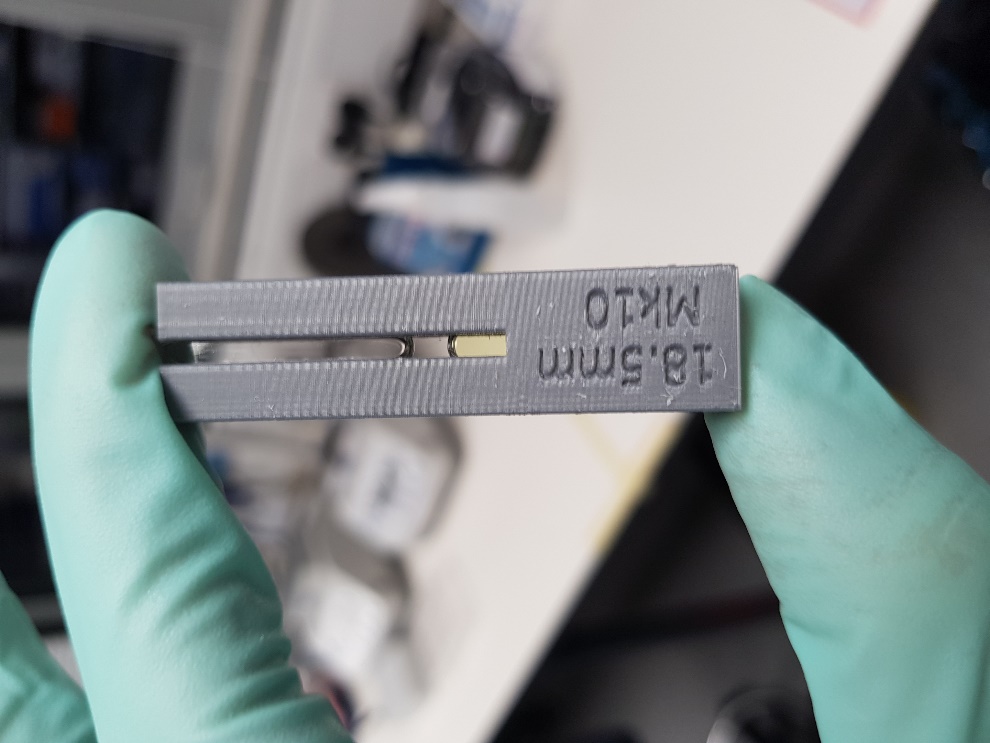

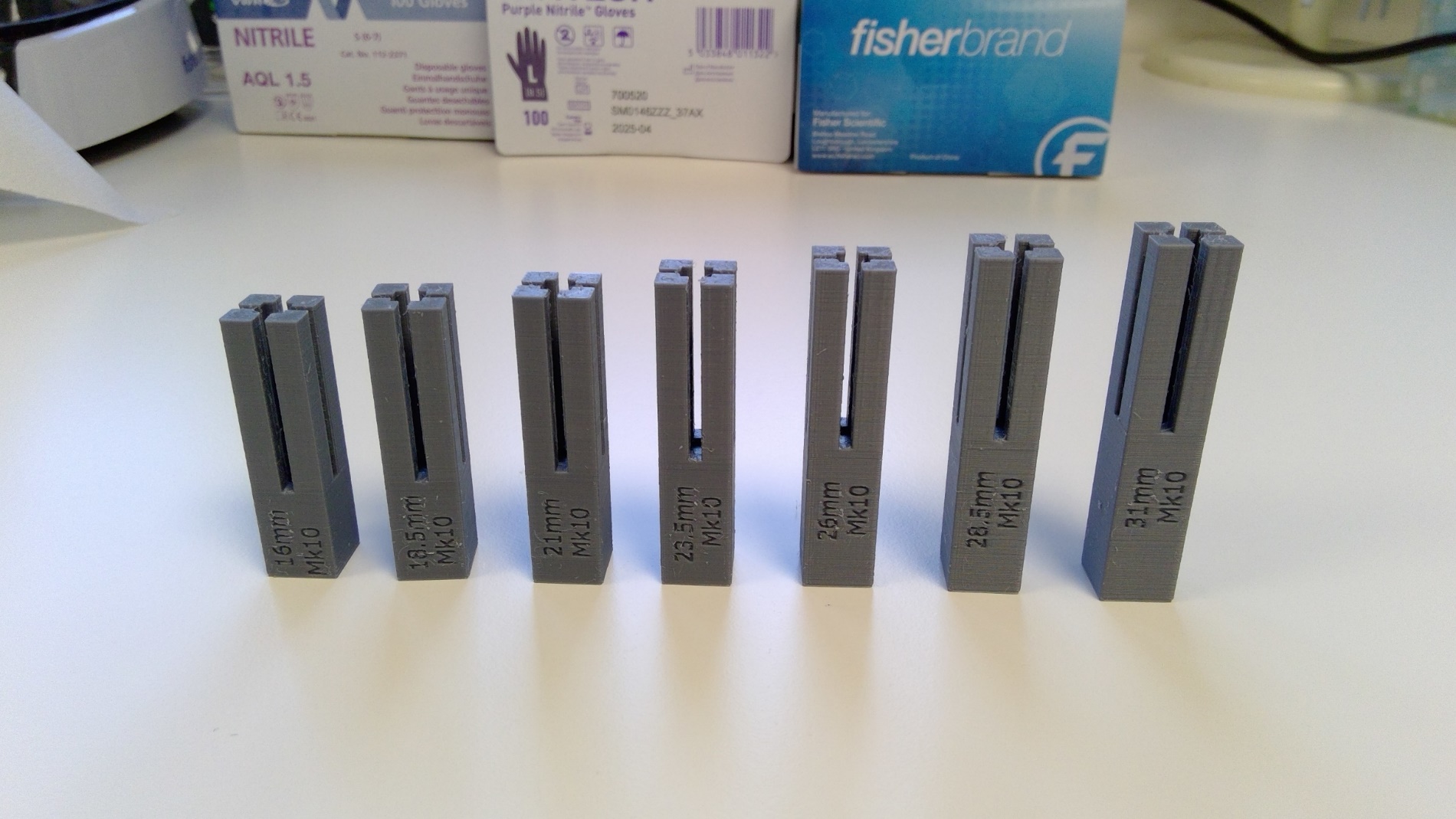

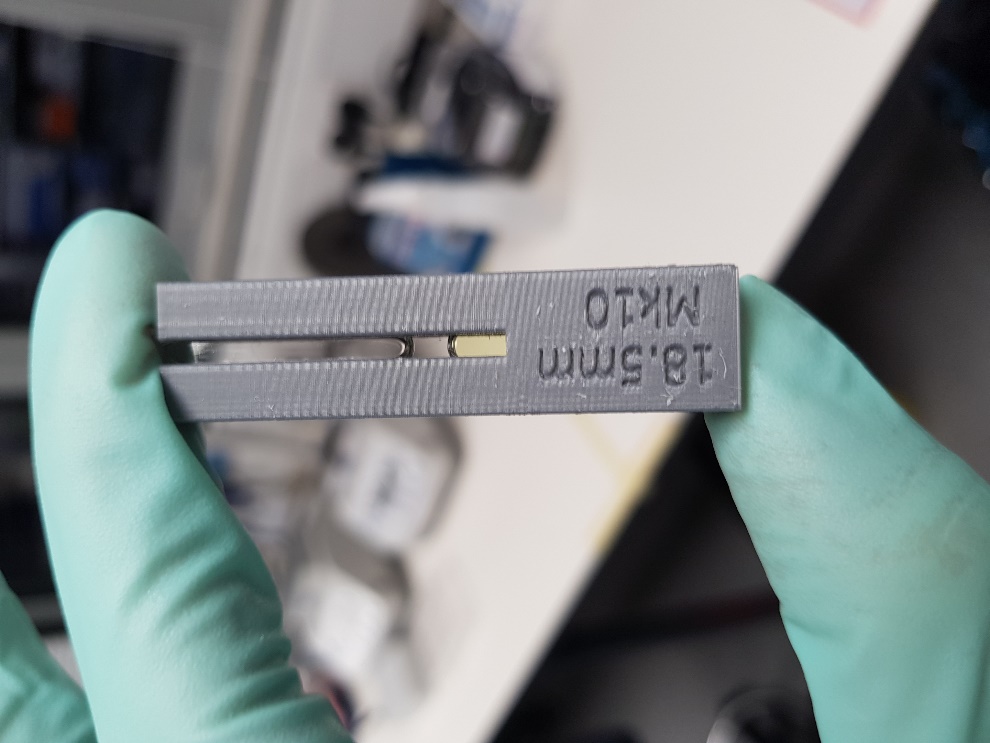

Microcuvette adapters were designed using Autodesk Inventor (v2019.1.1). 18.5mm was found to be the optimal height for the Agilent Cary Eclipse light path. Parts were printed on a Prusa Mk3S using PLA, 0.15mm layer height. Above image shows 100µL reaction mixture + 50µL mineral oil to prevent evaporation.
